# Model connectivity: leveraging the power of encoding models to overcome the limitations of functional connectivity

**DOI:** 10.1101/2023.07.17.549356

**Authors:** Emily X. Meschke, Matteo Visconti di Oleggio Castello, Tom Dupré la Tour, Jack L. Gallant

**Author notes:** These authors contributed equally (Emily X. Meschke, Matteo Visconti di Oleggio Castello).

## Abstract

Functional connectivity (FC) is the most popular method for recovering functional networks of brain areas with fMRI. However, because FC is defined as temporal correlations in brain activity, FC networks are confounded by noise and lack a precise functional role. To overcome these limitations, we developed model connectivity (MC). MC is defined as similarities in encoding model weights, which quantify reliable functional activity in terms of interpretable stimulus- or task-related features. To compare FC and MC, both methods were applied to a naturalistic story listening dataset. FC recovered spatially broad networks that are confounded by noise, and that lack a clear role during natural language comprehension. By contrast, MC recovered spatially localized networks that are robust to noise, and that represent distinct categories of semantic concepts. Thus, MC is a powerful data-driven approach for recovering and interpreting the functional networks that support complex cognitive processes.

## Introduction

Complex cognitive functions are subserved by anatomically and functionally coupled networks of brain areas^1–4^. Functional connectivity (FC) is the most popular method for recovering such networks with functional magnetic resonance imaging (fMRI)^5^. However, despite its popularity, FC suffers from two fundamental flaws that limit the reliability and interpretability of the recovered networks.

First, FC assumes that brain areas are functionally coupled if their blood-oxygenation-level-dependent (BOLD) activity is temporally correlated^6–9^. However, brain activity recorded from two brain areas can be temporally correlated even if the two brain areas do not share functional information. For instance, spurious correlations between two brain areas can arise from temporally correlated noise generated by physiological sources (e.g., respiration, heart-rate, movement^10, 11^). The magnitude of the spurious correlations between brain areas is affected by the signal-to-noise-ratio (SNR) of the brain areas^12, 13^. Two areas with low SNR, for instance, are more likely to exhibit spurious correlations resulting from temporally correlated noise. Because SNR varies across the brain (i.e., according to the participant’s head and brain anatomy, the participant’s vasculature, the hemodynamic response function, and the fMRI pulse sequence^11, 14–17^), the magnitude of the spurious correlations also varies across the brain. Therefore, FC is unavoidably confounded by spatial variations in SNR.

Second, FC does not provide a direct or precise method for assigning functions to the recovered networks. Most FC studies use a resting state paradigm, making it impossible to infer what functional information is reflected in the brain activity^18, 19^. However, even when studies use task-based paradigms such as movie watching^20–24^, FC does not provide any means to define the stimulus- or task-related information that accounts for the temporally correlated brain activity. Thus, most FC studies must infer function by comparing the cortical distribution of the recovered networks to maps of task activation^25–31^, or to networks that were functionally defined in previous studies^32, 33^. This approach is both inefficient, because it requires additional data or experiments, and imprecise, because it produces coarse functional descriptions that are associated with broad cognitive or sensory domains (e.g., “attention”, “motor”, or “visual”). Furthermore, this approach implicitly assumes that each network performs a single function across tasks, which is likely too strong of an assumption^34, 35^.

To overcome the limitations of FC, we developed a new method called model connectivity (MC). MC leverages the power of the Voxelwise Modeling (VM) framework, which builds encoding models for each voxel with features extracted from an experiment^36–42^. These features quantify the cognitive, sensory, or motor information that is hypothesized to be represented in the brain activity during the experiment. Regularized linear regression is used to predict brain activity from these features^43, 44^. Because the model weights are estimated through cross-validation and are evaluated on a separate test set, the model weights reflect reliable functional signals in the BOLD activity that generalize beyond the training set.

Because the model weights indicate which cognitive, perceptual, or motor features are represented in the brain activity, the model weights quantify the stimulus- or task-related functional tuning of individual voxels. MC uses these model weights to quantify functional coupling between brain areas, identifying networks of areas with similar functional tuning.

Note that we named our new encoding-model-based approach for recovering functional networks “model *connectivity*” to be consistent with common terminology. However, despite what their names suggest, neither functional connectivity nor model connectivity necessarily reflect true anatomical connectivity between brain regions^19, 45, 46^. Rather, functional connectivity captures patterns of temporal correlations in BOLD activity, and model connectivity captures patterns of functional similarity in terms of the stimulus- or task-related features.

To demonstrate the advantages of MC over FC, both methods were applied to a naturalistic fMRI dataset that was collected as participants listened to narrative stories^40^. We reasoned that the networks recovered from this narrative auditory dataset will reveal the functional organization of semantic processing systems in the brain^47^. For both MC and FC, the optimal sets of networks were identified by evaluating the cross-participant prediction accuracy of all possible sets of networks. To assess how well each of the MC and FC networks reflect the underlying functional activity, the cortical distribution, prediction accuracy, and functional signal quality were compared between MC and FC networks. To estimate the function of each FC network, the cortical distribution of the network was compared to the set of canonical resting state networks recovered by Yeo et al. (2011)^32^. To interpret the function of each MC network, the network-average functional tuning provided by the encoding model weights were manually inspected. Finally, the functional tuning and spatial organization of each MC network was assessed in each individual participant.

## Results

### Recovering functional networks with model connectivity and functional connectivity

To compare the networks recovered by model connectivity (MC) and functional connectivity (FC), both methods were applied to a naturalistic fMRI dataset (**Figure 1**). In this dataset, BOLD responses were recorded from 11 participants as they listened to narrative auditory stories^40^. Data from nine participants were used to recover the MC and FC networks reported here, and data from two participants were held-out for further validation of the networks (see **Methods**). For MC, encoding models were fit to each participant’s functional data. While the VM framework was developed to estimate functional tuning in each voxel in each participant’s native anatomical space, MC was developed to recover a single set of networks that reflect patterns of functional coupling that generalize across participants. Therefore, BOLD responses from each participant were first mapped to the fsaverage5 template surface, which consists of 10,242 vertices in each hemisphere (20,484 vertices total). Then, each participants’ functional data were modeled in terms of low-level auditory features (i.e., word rate, phoneme rate, and phonemes) and high-level semantic features (i.e., word co-occurrence rates) that had been extracted from the stories. Regularized linear regression was used to estimate the participant’s auditory and semantic model weights for each vertex of the fsaverage5 surface. Because these models separately account for the auditory and semantic information represented in the brain activity, the semantic model weights reflect the pure semantic tuning of each vertex in each participant, and are not confounded by low-level auditory features. These semantic model weights were then averaged across participants (see **Supplemental Figure 1** for prediction accuracy of the semantic encoding models in each participant), and MC was evaluated as the pairwise Euclidean distance between the group-average semantic model weights in all 20,484 vertices. For FC, BOLD responses that had been mapped to the template surface were averaged across participants, and FC was evaluated as the correlation between the group-average BOLD responses in all 20,484 vertices. To group vertices into networks, hierarchical clustering with Ward’s linkage was applied to the resulting MC and FC connectivity matrices.

**Figure 1.**
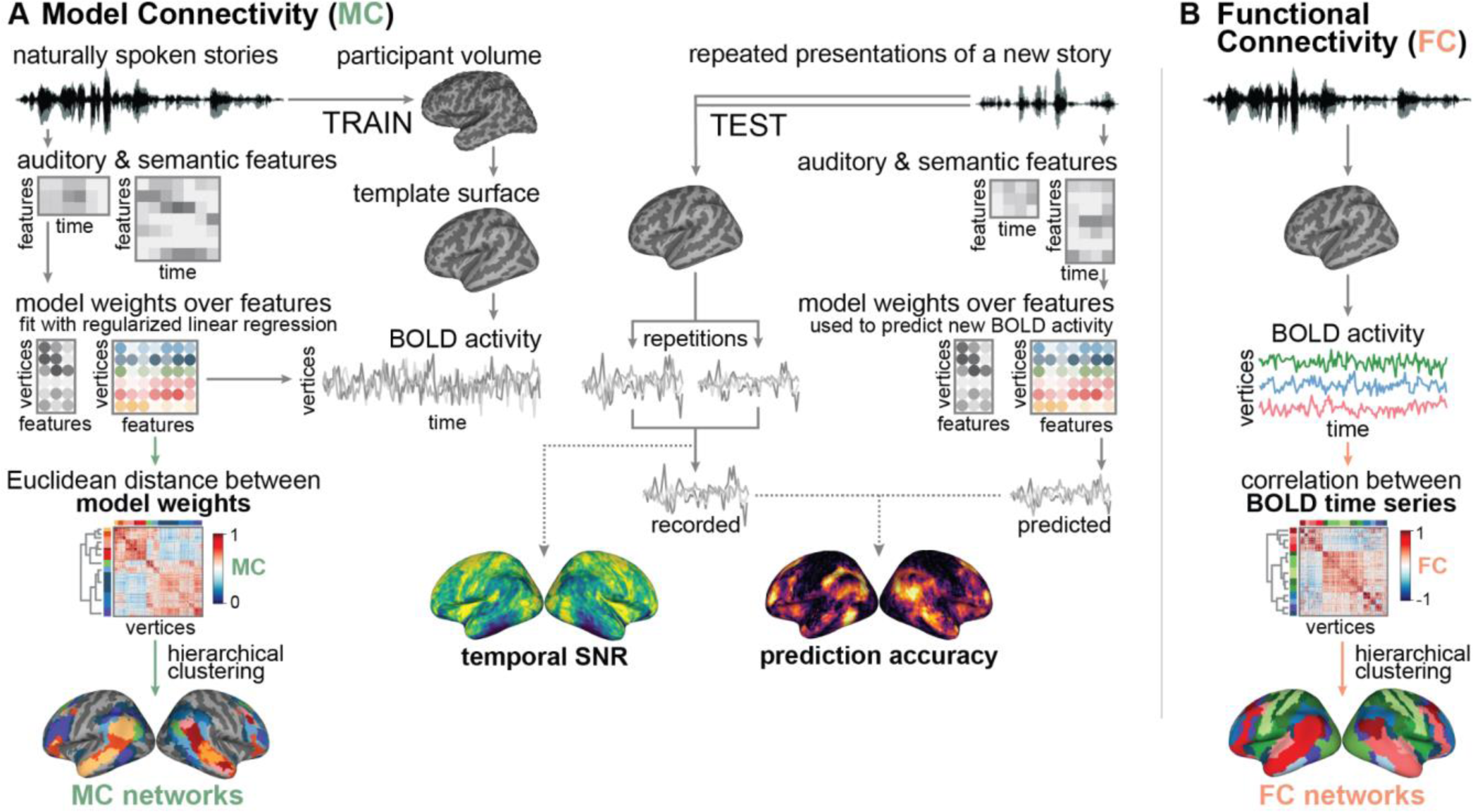
Model connectivity recovers functional networks from encoding model weights estimated using the Voxelwise Modeling framework (VM). (**A**) Model connectivity builds on VM to recover functional networks. In VM, stimulus- or task-related features are used to fit an encoding model that predicts brain activity in each voxel of each participant. The resulting encoding model weights provide a profile of the stimulus- or task-related functional tuning for each voxel in each participant. The quality of the functional data (temporal SNR) and the predictive power of the encoding models (prediction accuracy) are evaluated in a separate test set. MC uses the Euclidean distance between model weights to recover functional networks. In the present study, low-level auditory and high-level semantic features were extracted from narrative auditory stories^40^. Each participant’s brain activity in response to the stories was mapped from voxels in the participant’s native space to vertices of the template fsaverage5 surface. Brain activity in each vertex was modeled as a linear combination of the auditory and semantic features, and MC was applied to the semantic model weights. The resulting networks consist of vertices with similar semantic tuning. (**B**) In contrast, functional connectivity does not model brain activity. Rather, FC uses the correlation between BOLD time-series to recover functional networks. In the present study, brain activity in response to the narrative auditory stories was mapped to the template fsaverage5 surface, and FC was applied to these BOLD time-series. The resulting networks consist of vertices with similar temporal profiles of BOLD activity.

### Using encoding models to identify the optimal number of networks

One of the major challenges of recovering functional networks is determining the number of networks that underlie the observed patterns of functional coupling. FC studies typically use within-cluster homogeneity metrics that quantify the consistency of brain responses within the recovered networks^48–51^, or they use reliability metrics that quantify the stability of clustering solutions across participants^32, 52, 53^. However, spatial variability in SNR affects both the consistency of brain activity between brain regions and the consistency of brain activity across datasets^35^. Therefore, neither method ensures that the optimal set of networks accurately reflects the underlying functional organization rather than variability in noise levels across the brain.

To determine the set of networks that best captures the functional organization of the brain during the task, we developed a new method that leverages the encoding models fit to each participant’s functional data. Using a leave-one-participant-out cross-validation scheme, each clustering solution is evaluated in terms of how well the solution predicts brain activity in the test dataset of each left-out participant (**Figure 2A**). Cross-participant model prediction accuracy quantifies how well the encoding model weights averaged across the *N*-1 group of participants predict the stimulus- or task-related information reflected in the left-out participant’s brain activity. The optimal number of networks is defined as the number that maximizes the model prediction accuracy across left-out participants. Therefore, with this approach, the optimal number of networks produces the clustering solution that best captures the stimulus- or task-related patterns of functional coupling that are shared across participants. Note that because this approach relies on encoding models, it cannot be used by traditional FC studies. However, the aim of the present work is to compare MC and FC, so the semantic encoding model weights were used to evaluate both the MC and FC clustering solutions.

**Figure 2.**
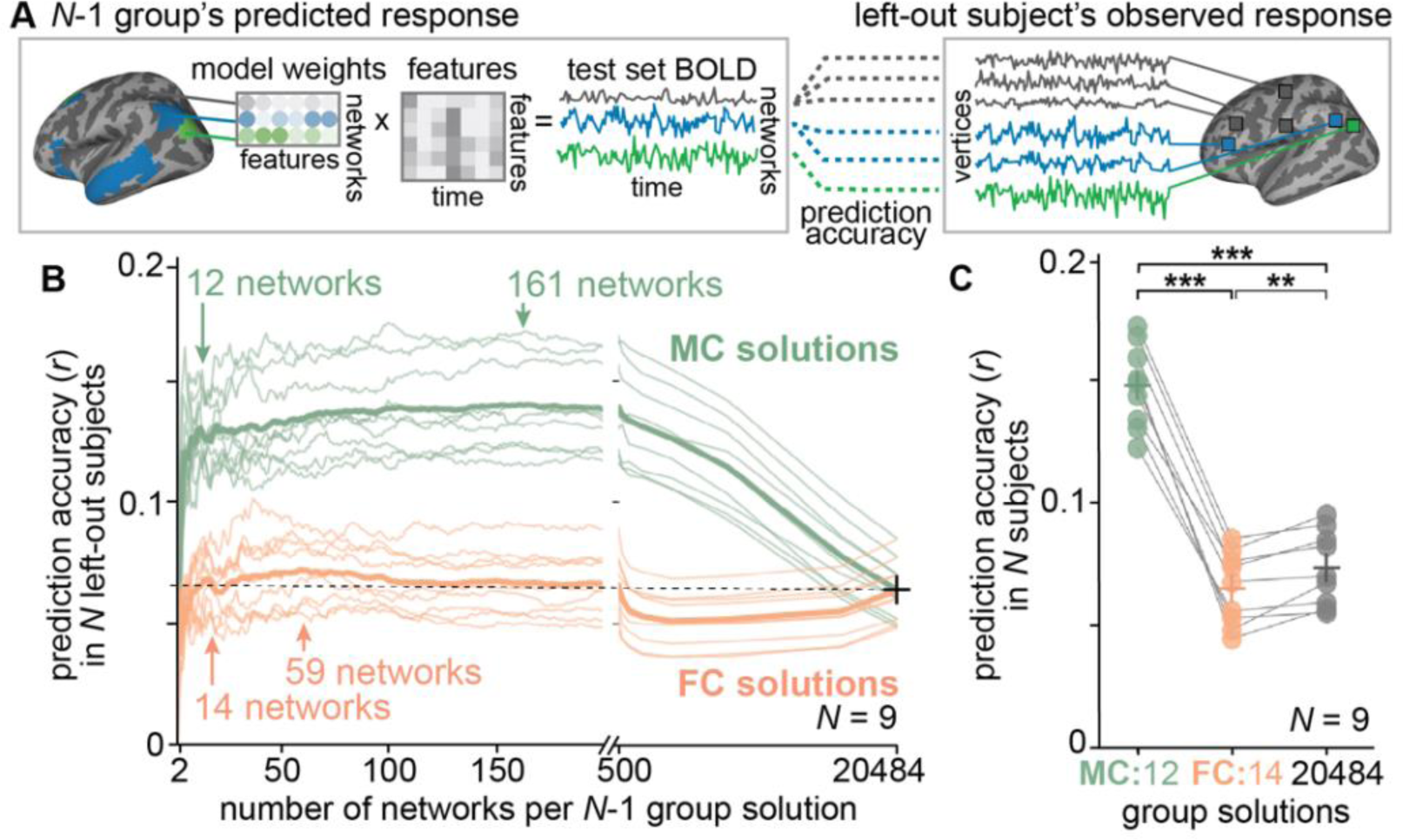
Encoding models are used to determine the number of networks that maximizes prediction accuracy across held-out participants. A leave-one-participant-out cross-validation scheme was used to identify the number of networks recovered by MC and FC that best predicted BOLD responses across participants. For each held-out participant, the remaining *N*-1 participants were used to generate the *N*-1 group solutions (*N* = 9). (**A**) To compute the prediction accuracy of each network in each clustering solution, the model weights from *N*-1 participants were first averaged across all vertices within the network, and were then used to predict BOLD responses in the test set of the held-out participant. Prediction accuracy of the network was computed as the average correlation between the network-average predicted and the vertex-wise observed BOLD response. Prediction accuracy of the clustering solution was computed as the average prediction accuracy across networks in the solution. (**B**) The prediction accuracies of all possible *N*-1 group solutions (y-axis) are plotted as a function of the number of networks in the solution (x-axis). MC solutions are plotted in green, and FC solutions are plotted in orange. Thin traces indicate the prediction accuracy for each held-out participant, and thick traces indicate the average prediction accuracy across held-out participants. The horizontal dotted line denotes the average prediction accuracy across all 20,484 vertices of the template fsaverage5 surface. Both MC and FC solutions show an early, sharp rise in prediction accuracy as the number of networks increases from 2 to 12 for MC, and from 2 to 14 for FC. Prediction accuracy continues to rise until the number of networks reaches 161 networks for MC and 59 networks for FC. To facilitate interpretation, the early local maxima of 12 and 14 were selected as the optimal number of networks. (**C**) The final MC and FC solutions were recovered using data from all 9 participants. The average prediction accuracy of the 12-network MC solution, 14-network FC solution, and 20,484-vertex solution are plotted for each participant. A line connects the prediction accuracy for each participant across the three solutions. The crosses indicate the average prediction accuracies across participants. Both the 12-network MC solution and the 20,484-vertex solution are significantly better predictors of BOLD responses across held-out participants than the 14-network FC solution (*** indicates p < 0.001, ** indicates p < 0.01, paired t-test). This indicates that MC, but not FC, groups vertices into networks that improve the cross-participant generalizability of the vertex-wise encoding models.

To identify the optimal number of networks, we examined the prediction accuracy of all possible MC and FC solutions, ranging from 2 networks to 20,484 networks (**Figure 2B**). For MC, prediction accuracy rises sharply as the number of networks per clustering solution increases from 2 networks to 12 networks (from *r* = 0.06 ± 0.02 to *r* = 0.13 ± 0.01, mean ± standard deviation (s.d.) across participants). Prediction accuracy continues to rise until the global maximum of 161 networks (*r* = 0.14 ± 0.02), and then declines as the number of networks approaches the maximum number of 20,484 networks (*r* = 0.06 ± 0.01). This decline in prediction accuracy reflects an overfitting to the group-average data, such that the group solutions with a large number of networks are too detailed to generalize to individual, left-out participants. For FC, prediction accuracy rises somewhat sharply as the number of networks per clustering solution increases from 2 networks to 10 networks (from *r* = 0.04 ± 0.02 to *r* = 0.07 ± 0.01). Prediction accuracy continues to rise slightly until the global maximum of 59 networks (*r* = 0.07 ± 0.01), and then declines slightly as the number of networks approaches the maximum number of 20,484 networks. Thus, for MC, the cross-participant prediction accuracy is maximized at 161 networks, and for FC, the cross-participant prediction accuracy is maximized at 59 networks (see **Supplemental Figure 2** for the 161-network MC solution and the 59-network FC solution). These large numbers of networks are the most accurate reflection of the functional subdivisions that can be recovered by MC and FC from this dataset. However, for the sake of exposition in the present work, we focused all subsequent analyses on the set of 12 networks recovered by MC and the set of 14 networks recovered by FC.

To recover the final sets of 12 MC and 14 FC networks, we used data from the entire group of *N* participants. Here we refer to the resulting solutions as the group 12-network MC solution and the group 14-network FC solution. These clustering solutions are called the *group solutions* to distinguish them from the MC and FC clustering solutions that were generated using different sets of *N*-1 participants in the cross-validation scheme. In the remainder of this paper, all references to the 12-network MC solution and the 14-network FC solution refer to these two group solutions that were recovered from the entire group of *N* participants. The cross-participant prediction accuracy was then computed for the final MC and FC group solutions, to assess how well these solutions predict the underlying brain activity in individual participants (**Figure 2C**). For each participant, the 12-network MC solution was significantly better at predicting the participant’s BOLD responses in a held-out test set than the 14-network FC solution (difference = 0.08, *t*(8) = 20.53, *p* < 0.001, paired t-test). This indicates that the 12 networks recovered by MC more accurately reflect the organization of the functional networks that support natural language comprehension.

The MC and FC solutions can be viewed as low-dimensional reconstructions of the vertex-wise functional map, in which the full-resolution map comprising 20,484 vertices is reduced to 12 MC networks and 14 FC networks respectively. Because the dimensionality of the reconstruction is determined according to the cross-participant prediction accuracy, we expected the optimal dimensionality to be a trade-off between functional specificity and functional generalizability. While the 12-network MC solution and the 14-network FC solution do not reflect the fine-grained differences between individual participants, they may recover the coarse functional organization that is shared between participants, and therefore may better generalize across participants.

To test whether the 12-network MC solution or the 14-network FC solution improves the cross-participant generalizability of the vertex-wise encoding models, the average prediction accuracy across networks in the MC and FC solutions was compared to the average prediction accuracy across all 20,484 vertices of the fsaverage5 template surface (**Figure 2C**). Grouping the 20,484 vertices of the fsaverage5 template surface into the 12 MC networks significantly improves the prediction accuracy in each individual participant (difference = 0.07, *t*(8) = 18.64, *p* < 0.001, paired t-test). This indicates that reducing the full-resolution vertex-wise encoding models into the 12 MC networks improves the cross-participant generalizability of the vertex-wise encoding models. Therefore, these 12 MC networks are an effective, low-dimensional reconstruction of the functional organization that supports natural language comprehension and that is shared across individuals. In contrast, grouping the 20,484 vertices into the 14 FC networks significantly reduces the prediction accuracy in each individual participant (difference = −0.01, *t*(8) = −4.24, *p* < 0.01, paired t-test). The 14 FC networks therefore do not improve the cross-participant generalizability of the vertex-wise encoding models, and consequently do not effectively reflect the functional organization that supports natural language comprehension.

### Model connectivity recovers spatially localized networks that accurately reflect the functional organization related to the experimental task

The 12-network MC solution and the 14-network FC solution reflect two hypotheses about how the brain is functionally organized during natural language comprehension. The previous analysis revealed that the 12-network MC solution is a more effective low-dimensional reconstruction of the vertex-wise functional maps than the 14-network FC solution. This indicates that the functional organization recovered by the MC solution reflects the functional organization of language processing areas more closely than the functional organization recovered by the FC solution. We therefore compared the similarity structure, cortical distribution, and prediction accuracy of each network in the 12-network MC solution (**Figures 3A-C**) and in the 14-network FC solution (**Figures 3D-F**) to assess how the two solutions differ in terms of the recovered functional organization.

**Figure 3.**
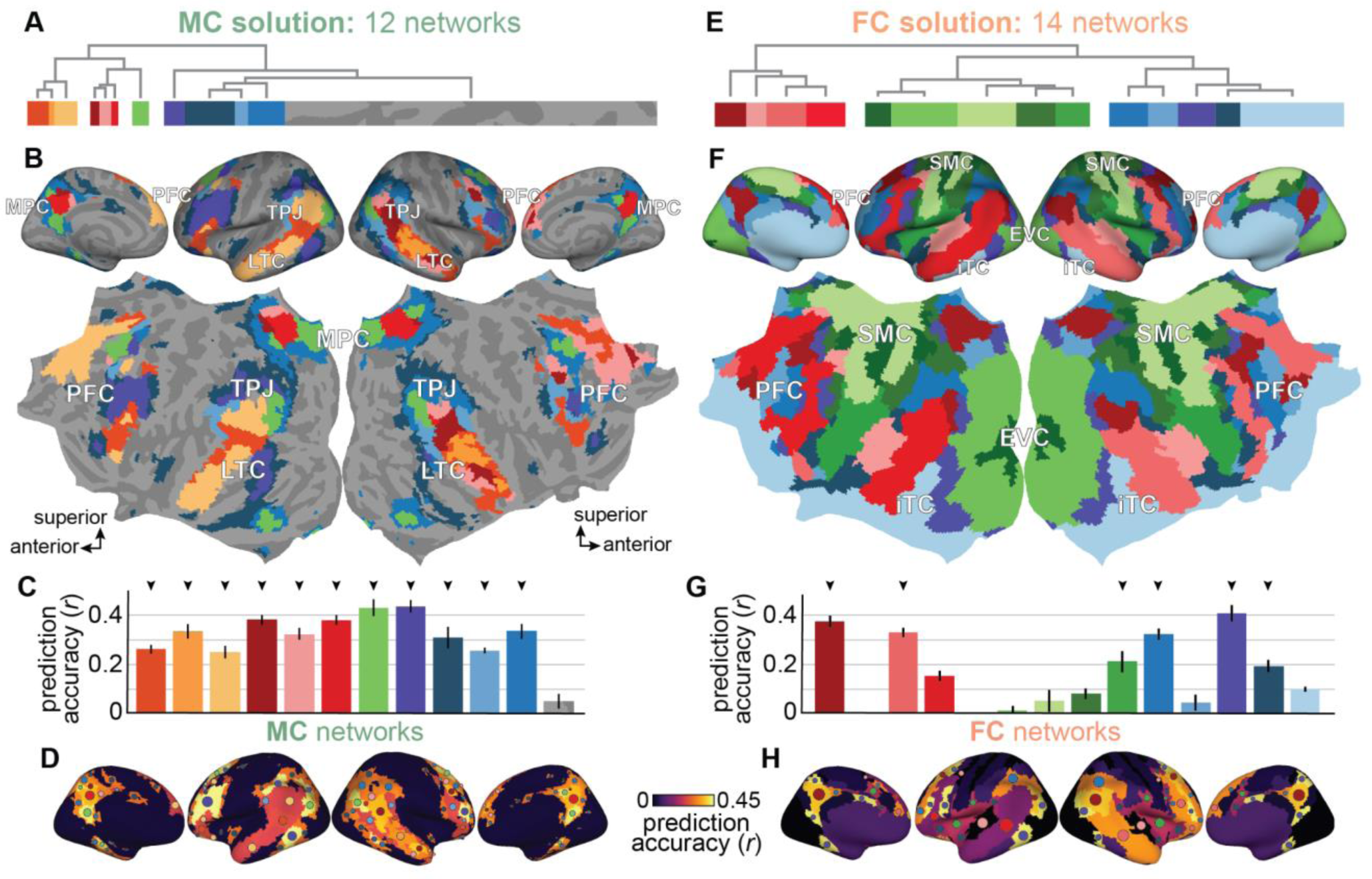
Model connectivity recovers spatially localized networks that accurately reflect the functional organization of the brain during natural language comprehension. The optimal MC and FC solutions reflect hypotheses about the functional organization of the brain during natural language comprehension. To compare the functional organization depicted by these two solutions, we evaluated the similarity structure, the cortical distribution, and the prediction accuracy of each network in the MC and FC solutions. (**A**, **E**) The dendrograms show the similarity across networks in (**A**) the 12-network MC solution and (**E**) the 14-network FC solution. The color of each bar indicates the network assignment, and the length of each bar corresponds to the number of vertices in the network. Visual inspection of the dendrograms revealed that the 12-network MC solution can be divided into four groups of networks, and the 14-network FC solution can be divided into three groups of networks. To preserve this high-level similarity structure, each group of networks was assigned a distinct color. Note that the largest network in the 12-network MC solution was not assigned a color, and is instead represented by patterns of gray cortical curvature. (**B**, **F**) The (**B**) MC and (**F**) FC solutions are visualized on the inflated and flattened template surface. Vertices are colored according to their network assignment, as in (**A**) and (**E**). In the MC solution, the 11 smaller networks span brain areas involved in processing the semantic information in natural language (TPJ, MPC, LTC, PFC). The largest network (represented by patterns of cortical curvature) spans areas that do not reliably respond to the semantic information in the narrative auditory stimulus (EVC, SMC), and areas that suffer from signal dropout (iTC). In the FC solution, five of the 14 networks span areas not involved in processing the semantic information in natural language (colored in shades of green), and one network spans areas with poor SNR (colored in light blue). Thus, while MC isolates all vertices that are not functionally relevant to the stimulus- or task-related features in a single network, FC recovers multiple networks that reflect activity that is not related to the stimulus or task. (**C**, **G**) The prediction accuracy of each network in the (**C**) MC and (**G**) FC solutions is plotted in a bar graph. The color of each bar corresponds to the network assignment, and the error bars indicate the standard error of the mean (s.e.m.) across the 9 participants. 11 of the 12 networks recovered by MC significantly predict brain activity in the majority of participants, whereas only six of the 14 networks recovered by FC significantly predict brain activity in the majority of participants (arrows indicate *p* < 0.05, permutation test). (**D**, **H**) The prediction accuracy of the (**D**) MC and (**H**) FC solutions is plotted on the inflated template surface. The colored circles indicate the significantly predicted networks, and are roughly scaled according to the size of each contiguous region within the network. While the significantly predicted networks of the MC and FC solutions span a similar range of cortical regions, MC recovers more functionally distinct subdivisions within these cortical regions. These analyses indicate that the 12-network MC solution is a more accurate and spatially localized depiction of the semantic systems that support language comprehension.

Because hierarchical clustering was used to group vertices into networks, the resulting dendrogram can be used to examine the similarity between networks in the MC and FC solutions. The dendrogram of the 12-network MC solution reveals that these 12 networks can be broadly divided into four groups (**Figure 3A**). Because the functional tuning within each group of networks is more similar than the functional tuning between groups of networks, each group of networks was assigned a distinct color. Thus, the 12-network MC solution consists of three networks colored in shades of orange, four networks colored in shades of red, one network colored in green, and four networks colored in shades of blue. Note that because one network stood out as being significantly larger than the other 11 networks (12,944 vertices compared to 685 ± 448 vertices, mean ± s.d.; *t*(11) = 86.59, *p* < 0.001, 1-sample t-test), this network was not assigned a color, and is instead represented by patterns of cortical curvature in gray. The dendrogram of the 14-network FC solution reveals that these 14 networks can be broadly divided into three groups according to the similarity of their BOLD responses (**Figure 3E**). Thus, the 14-network FC solution consists of four networks colored in shades of red, five networks colored in shades of green, and five networks colored in shades of blue. While the color of the networks reflects the similarity structure within the MC and FC solutions, the colors are not meaningfully matched between the two solutions.

Of the 12 networks recovered by MC (**Figure 3B**), 11 span brain areas known to process the semantic information in natural language (e.g., temporal parietal junction (TPJ), lateral temporal cortex (LTC), medial parietal cortex (MPC), prefrontal cortex (PFC)). The remaining network is the largest network, spanning brain areas that were not reliably activated by the semantic information in the narrative stories (e.g., somatomotor cortex (SMC) and early visual cortex (EVC)), and areas with low SNR (e.g., inferior temporal cortex (ITC)). Thus, the 12-network MC solution appears to comprise 11 networks that accurately reflect functional activity related to semantic information in natural language, and a single large network that isolates vertices that are not functionally involved in processing semantic information in natural language. In contrast, of the 14 networks recovered by FC (**Figure 3F**), six networks span a similar set of brain areas that are known to process language (the networks colored in dark red, pink, red, blue, medium blue, and dark purple). Of the eight remaining FC networks, one network spans early auditory cortices (colored in light pink), one spans insular and posterior prefrontal regions (colored in dark blue), five span somatomotor and early visual cortices (colored in shades of green), and one spans inferior areas that suffer from signal dropout (colored in light blue). Thus, given the cortical distribution of the recovered networks, the 14-network FC solution appears to recover six networks that are not involved in the task (colored in shades of green and in light blue). In sum, the MC solution separates brain areas that encode stimulus- or task-related information from those that are stimulus- and task-irrelevant, while the FC solution includes all areas regardless of their functional relevance to the stimulus or task.

The relative sizes and cortical distributions of the MC networks suggest that MC recovers networks that solely reflect the specific stimulus- or task-related patterns of functional coupling that are captured by the encoding models. To quantify the functional relevance of each MC network, we evaluated how well the group-average semantic model weights within each network predict each participant’s BOLD responses during the task (**Figure 3C**). The semantic model weights within 11 of the 12 MC networks significantly predict BOLD responses in at least seven of the nine participants (*p* < 0.05, permutation test; see **Supplemental Figure 3** for the prediction accuracy of the group-average semantic models in each network of each participant). The semantic model weights within the remaining MC network significantly predict brain activity in only one participant. Thus, as expected given its topographic organization, the 12-network MC solution consists of 11 networks that include areas that represent semantic information in natural language, and one noise network that spans all remaining brain areas that do not represent semantic information. The 12-network MC solution therefore defines 11 networks that reflect the precise functional organization of semantic processing systems in the brain. In the remainder of the paper, these 11 networks will be referred to as the 11 *semantic* networks.

The relative sizes and cortical distributions of the FC networks suggest that FC recovers patterns of functional coupling that reflect both functionally relevant and functionally irrelevant fluctuations in BOLD activity. Because FC does not use encoding model weights to recover functional networks, traditional FC studies cannot use prediction accuracy to evaluate the functional relevance of the networks. However, because the aim of the present work is to directly compare the networks recovered by MC and FC, we used the group-average semantic model weights estimated for MC to evaluate the functional relevance of each FC network (**Figure 3G**). The semantic model weights in six of the 14 FC networks significantly predict brain activity in at least seven of the nine participants (indicated by arrows). The semantic model weights within the eight remaining FC networks significantly predict brain activity in fewer than half of the nine participants. Thus, the 14-network FC solution defines six networks that reflect the functional organization of semantic processing systems in the brain, and eight networks that reflect patterns of temporal correlations that do not have a clear functional role.

To assess whether a similar range of cortical regions were well predicted by the MC and FC solutions, the network-average prediction accuracy of each network was mapped onto the inflated template surface (**Figures 3D** and **3H**). The MC and FC networks that significantly predict brain activity across left-out participants span a similar range of cortical regions that are known to be involved in processing semantic information in speech (Dice coefficient between these six FC networks and the 11 semantic MC networks = 0.79). However, the 12-network MC solution recovers 11 functionally distinct networks within these regions, whereas the 14-network FC solution recovers only six functionally distinct networks within these regions. Thus, the 12-network MC solution is a more spatially localized depiction of semantic representations across the brain.

### Model connectivity is not confounded by differences in temporal SNR across the brain

The previous analysis revealed that MC separates regions that are functionally tuned to the stimulus- or task-related features from regions that are functionally irrelevant, isolating these functionally irrelevant regions into a single “noise” network. By contrast, FC recovers several networks that are not functionally relevant to the stimulus or task. This suggests that while MC can correctly separate signal and noise in the measured BOLD responses, FC is confounded by noise in the measured BOLD responses. To test whether MC is less susceptible to noise than FC, we quantified the degree to which the MC and FC solutions capture spatial variability in SNR. To measure SNR, we used temporal SNR (tSNR). Temporal SNR quantifies the stability of the BOLD response over time, and is commonly used to assess the signal quality of fMRI data^54, 55^. A single group-average tSNR map was obtained by first computing a separate tSNR map for each participant, and then averaging tSNR maps across participants. Network-average tSNR maps were computed for the MC and FC solutions by averaging the vertex-wise tSNR within each network of each solution.

To quantify the degree to which the MC and FC solutions are confounded by spatial variability in tSNR, we computed the correlation between the vertex-wise tSNR maps and the network-average tSNR maps. A high correlation indicates that vertices with a similar tSNR are grouped together in the same network, and that the clustering solution is confounded by spatial variability in SNR. A correlation that is close to zero indicates that vertices with similar tSNR are not grouped together, and that the solution is not biased by differences in SNR across the brain. Significance was separately evaluated for the MC and FC solutions by comparing the observed correlation with a null distribution. For each solution, the null distribution was generated by first randomly rotating the solution around the inflated template surface, and then correlating the vertex-wise tSNR with the network-average tSNR of the randomly rotated solution (see **Methods** for more details).

The vertex-wise tSNR (**Figure 4A**) was compared to the network-average tSNR of the 12-network MC solution (**Figures 4B** and **C**) and to the network-average tSNR of the 14-network FC solution (**Figures 4D** and **E**). For the MC solution, the correlation between network-average and vertex-wise tSNR is not significantly higher than the null distribution (*r* = 0.08, *p* = 0.996, permutation test), indicating that the 12-network MC solution accounts for an insignificant portion of the spatial variance in tSNR. For the FC solution, the correlation between the network-average and vertex-wise tSNR is significantly higher than the null distribution (*r* = 0.63, *p* < 0.001), indicating that the 14-network FC solution accounts for a significant proportion of the spatial variance in tSNR (*r*^2^ = 0.40). Thus, while MC is robust to spatial variability in SNR, FC is confounded by differences in SNR across the cerebral cortex (see **Supplemental Figure 4** for the correspondence between the vertex-wise and network-average tSNR in each individual participant).

**Figure 4.**
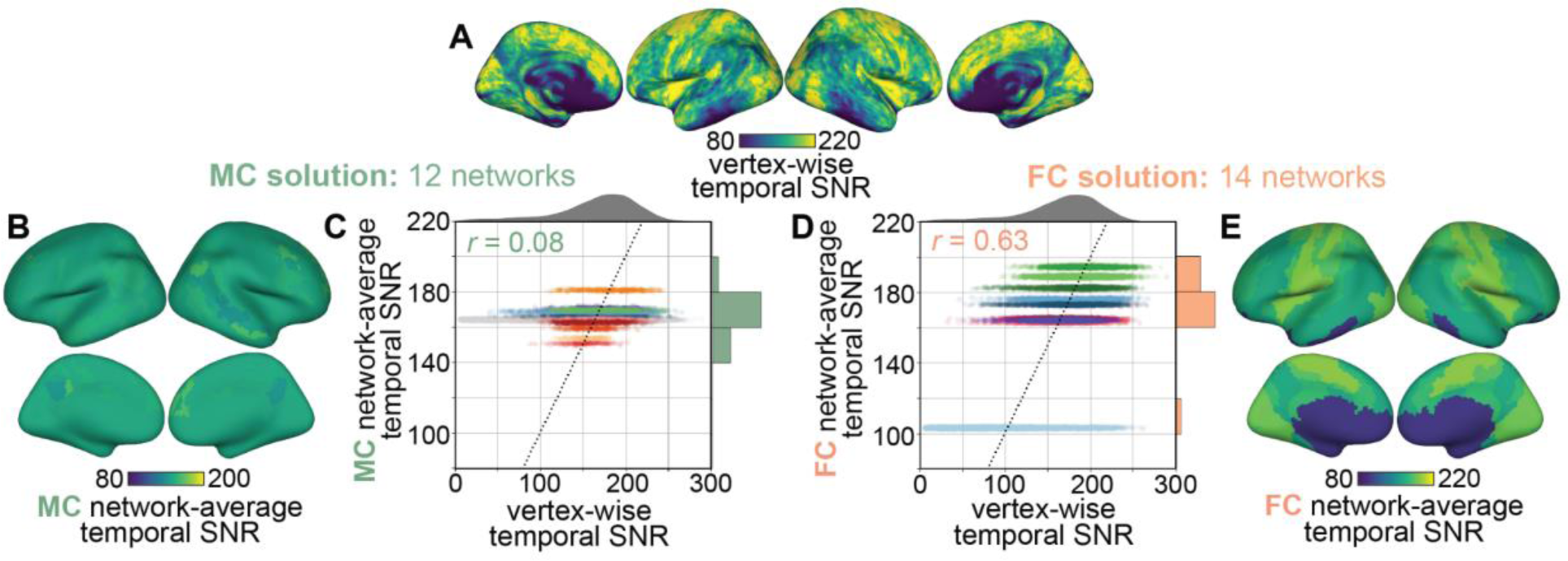
Model connectivity is not confounded by spatial variability in temporal SNR. Because FC is based on temporal correlations in BOLD activity, we hypothesized that variations in the signal quality of the recorded BOLD activity will affect the cortical distribution of the recovered networks. To test this hypothesis, we evaluated how closely the network-average temporal SNR (tSNR) reflects spatial variability in the vertex-wise tSNR. (**A**) For each vertex of the template surface, tSNR was first computed using functional data from each participant, and was then averaged across participants. The group vertex-wise tSNR is visualized on the inflated template surface. (**B**, **E**) The network-average tSNR is shown on the inflated template surface for each network in the (**B**) 12-network MC solution and (**E**) 14-network FC solution. Only the FC solution mirrors the spatial variability in tSNR shown in (**A**). (**C**, **D**) To quantify the degree to which the (**C**) MC and (**D**) FC solutions reflect the spatial pattern of tSNR, the vertex-wise tSNR was correlated with the network-average tSNR. In the scatter plot, each point represents a vertex, and the black dotted line indicates the identity line. The distribution of vertex-wise tSNR is shown above each scatter plot, and the distributions of network-average tSNR are shown to the right of each scatter plot. For the MC solution, the correlation between vertex-wise and network-average tSNR is near zero (*r* = 0.08). In contrast, for the FC solution, the correlation between vertex-wise tSNR and network-average tSNR is relatively high (*r* = 0.63). This indicates that while the networks recovered by MC are robust to spatial differences in SNR, the networks recovered by FC are confounded by spatial variability in SNR.

### Interpreting networks according to their cortical distribution results in coarse functional definitions

FC studies aim to recover the brain networks that support complex cognitive functions. However, in both task-based and resting-state FC studies, FC provides no means to define the functional information reflected in the BOLD responses. Consequently, the recovered networks do not have an explicit functional assignment. To circumvent this issue, FC studies define network functions in terms of broad functional correspondences with conventional task-based contrast studies. For example, if an FC network spans the occipital lobe, this network would be broadly defined as a visual network. The functional definitions resulting from this sort of analysis are necessarily coarse. Furthermore, because either the cortical distribution or the functional role of the networks recovered during one task may not generalize to another task, it may be misleading to use different tasks to recover networks and to define their functions.

To demonstrate the limitations of this traditional FC approach, we used the networks recovered and interpreted by Yeo et al.^32^ to assign a function to each MC and FC network recovered in the present work. Yeo et al. identified stable 7-network and 17-network solutions by applying functional connectivity to a large resting-state dataset. The function of each network in the 7-network solution was assigned by comparing the cortical distribution of each network to networks identified previously in the literature. This 7-network solution comprises the visual, somatomotor, limbic, ventral-attention, dorsal-attention, control, and default-mode network. The 17-network solution subdivides these 7 networks into two visual, two somatomotor, two limbic, two ventral-attention, two-dorsal attention, three control, one temporoparietal, and three default-mode networks (these functions are reported in Kong et al.^56^). The networks in the higher resolution 17-network solution are used here as templates for the canonical resting state functional networks (**Figure 5A**).

**Figure 5.**
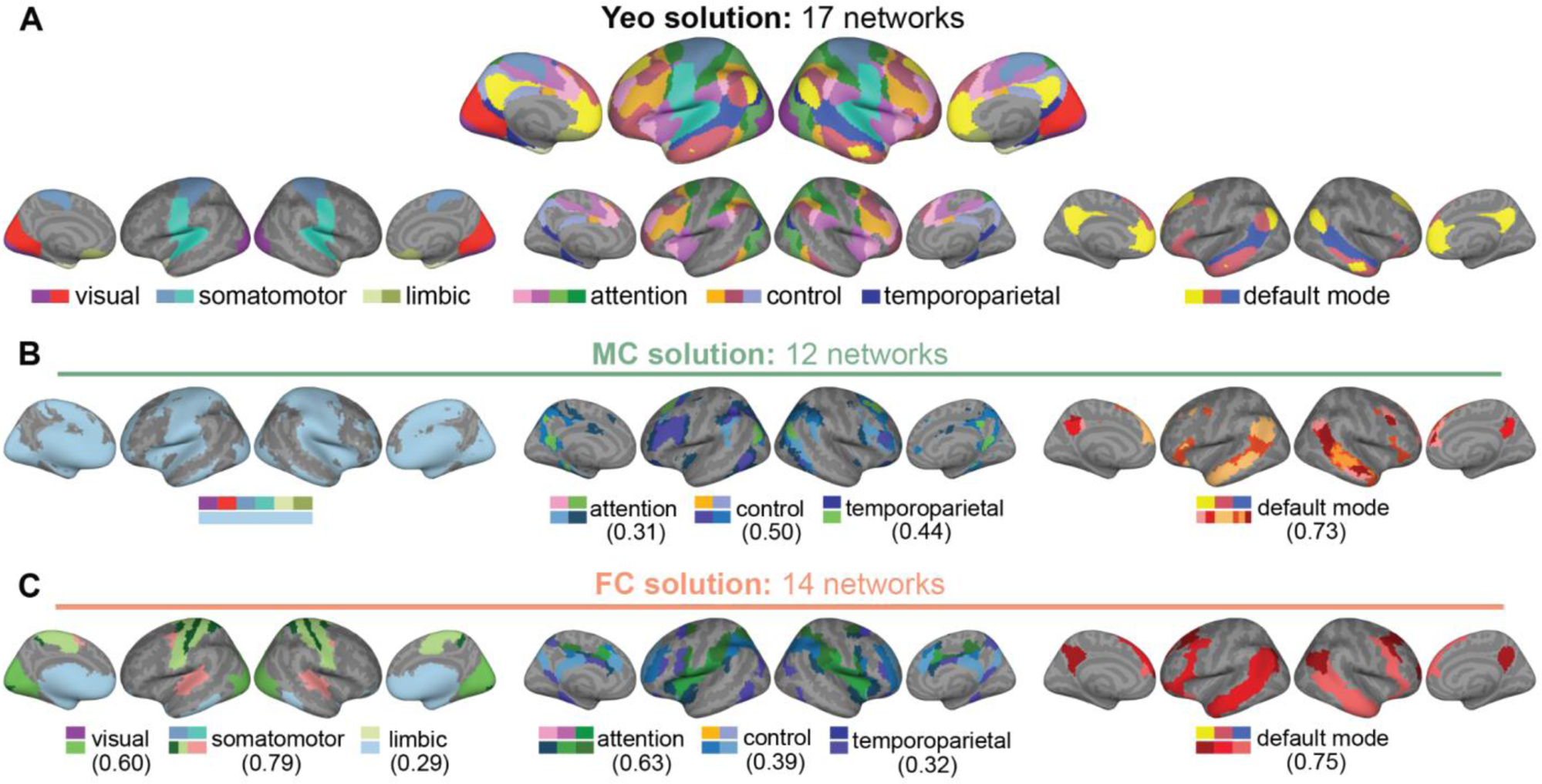
Relying on additional datasets to interpret the recovered networks results in coarse functional definitions that may not generalize across datasets. Because FC does not define the functional information reflected in the BOLD activity, traditional FC studies infer the function of the recovered networks by referencing what is already known about the functional organization of the brain. In the present work, we demonstrate the limitations of this approach by approximating the function of each MC and FC network as the function of one of the 17 resting state networks recovered by Yeo et al.^32^. (**A**) The 17-network Yeo solution is shown on the inflated template surface. These networks were recovered in a dataset of 1000 participants at rest, and reflect the set of canonical resting state functional networks. To facilitate visualization, the 17 networks were split into three groups (visual, somatomotor, and limbic; attention, control, and temporoparietal; default mode), and are shown separately on the inflated template surface. The broad function assigned to each network is denoted below the inflated surfaces. (**B**) For each network in the 12-network MC solution, the Dice coefficient was used to determine the best matching Yeo network. The MC networks were split into the same three groups as in (**A**), and are shown on the inflated surface. The colored squares indicate the best matches between the Yeo and MC networks. The Dice coefficient (reported in parentheses) quantifies the similarity in cortical distribution between the union of the Yeo networks and the union of the best matching MC networks. According to this approach, applying MC to a narrative auditory dataset recovers variations of the attention, control, temporoparietal, and default mode networks. The largest “noise” network comprises the remaining visual, somatomotor, and limbic networks that are not reliably activated by natural language comprehension. (**C**) For each network in the 14-network FC solution, the Dice coefficient was used to determine the best matching Yeo network, as in (**B**). According to this approach, applying FC to a narrative auditory dataset recovers variations of all canonical resting state networks. However, because this approach does not define the functional role of these networks during the task, these coarse functional definitions do not provide further insight into the functional organization of the brain during natural language comprehension.

In traditional FC studies, the function of the recovered network is assigned by visually inspecting the overlap between the recovered network and the template networks. Here, we used a less subjective approach that relies on the Dice coefficient to quantify the similarity between the cortical distribution of each Yeo network and each network recovered in the present work^50, 56, 57^. The Dice coefficient ranges from 0 to 1. A Dice coefficient of zero indicates that two networks do not share any vertices, and a Dice coefficient of one indicates that two networks share all vertices. For each MC and FC network, the 17 Yeo networks were ranked according to their Dice coefficient. The function of the MC or FC network was estimated to be the same as the Yeo network with the largest Dice coefficient. The MC and FC networks were then grouped according to their functional assignment, and the Dice coefficient between the union of each group of networks and the union of the corresponding resting state networks was computed. The resulting Dice coefficients quantify how well the canonical resting state networks can be recovered from the natural language comprehension task dataset (denoted below as *D*).

Using this approach, we determined that MC recovers variations of four of the seven (or eight of the 17) canonical resting state networks (**Figure 5B**). Specifically, two MC networks are variations of two of the four attention networks (*D* = 0.31, *p* < 0.001; permutation test), two MC networks are variations of two of the three frontoparietal control networks (*D* = 0.50, *p* < 0.001), one MC network matches the single temporoparietal network (*D* = 0.44, *p* < 0.001), and six MC networks subdivide the three default mode networks (DMN; *D* = 0.73, *p* < 0.001). The “noise” MC network spans all of the remaining resting state networks (i.e., the two visual networks, the two somatomotor networks, and the two limbic networks). By contrast, we found that FC recovers variations of all seven of the seven (or 13 of the 17) canonical resting state networks (**Figure 5C**). Specifically, one FC network is most similar to one of the two visual networks (*D* = 0.60, *p* < 0.001), three FC networks subdivide the two somatomotor networks (*D* = 0.79, *p* < 0.001), one FC network is best matched by one of the two limbic networks (*D* = 0.29, *p* < 0.01), three FC networks are variations of three of the four attention networks (*D* = 0.63, *p* < 0.001), two FC networks are best matched by two of the three frontoparietal control networks (*D* = 0.39, *p* < 0.001), one FC network matches the single temporoparietal network (*D* = 0.32, *p* < 0.001), and three FC networks are variations of the three DMNs (*D* = 0.75, *p* < 0.001). The results of this analysis are largely consistent with our previous finding that MC networks span functionally-relevant brain areas, while FC networks span all brain areas regardless of their functional relevance. Furthermore, these results suggest that regions associated with the canonical DMN are important for language processing^40^, as both the MC and FC solutions contain a set of networks that are well-matched to the DMN. However, because the functional definitions of all of these resting state networks are not specific to the experimental task, this standard approach for interpreting network functions does not provide further insight into the functional organization of the brain during natural language comprehension.

### Interpreting networks according to their functional tuning results in precise functional definitions that are relevant to the experimental task

While the previous analysis recovered coarse functional definitions for each MC and FC network, further attempts to interpret the networks recovered by FC are limited. With FC alone, there is not enough information to identify the factors that account for the observed differences in cortical distribution between the canonical resting state networks and the task-based networks. Furthermore, there is not enough information to identify the functional role of each of these canonical networks in the natural language comprehension task. In contrast, with MC, each network is associated with a high-dimensional functional tuning profile described by the encoding model weights. Therefore, the functional role of each of the recovered networks during the experimental task is well defined. In the present study, we used the semantic encoding model weights to recover 11 semantic networks and to identify the categories of semantic concepts represented by each network. First, the semantic model weights were averaged across vertices within each network. Then, the network-average model weights were used to identify the words spoken in the stories that are predicted to elicit the largest BOLD response in the network. Lastly, these words were manually inspected to interpret the precise concepts represented by the network.

To aid visualization, the 11 semantic networks were split into four groups according to the similarity of their semantic tuning (**Figure 6A**). To assess the degree to which each network represents a distinct category of semantic concepts, the 100 words (the top ∼1% of the 10,470 words spoken in the stories) that were predicted to elicit the largest BOLD response in each network were plotted on a two-dimensional projection of the high-dimensional semantic space (**Figure 6B**). In the two-dimensional semantic space, semantically similar words are located close to each other. Therefore, for each network, the distribution of the top 100 words across this reduced semantic space indicates the breadth of semantic concepts represented by the network. Visually inspecting the spatial distributions of these words for each network revealed that seven of the 11 semantic networks represent distinct semantic categories. The four remaining networks can be grouped into two pairs of networks, such that each pair of networks represents a similar range of semantic concepts. Thus, the 11 semantic networks represent nine distinct categories of semantic concepts.

**Figure 6.**
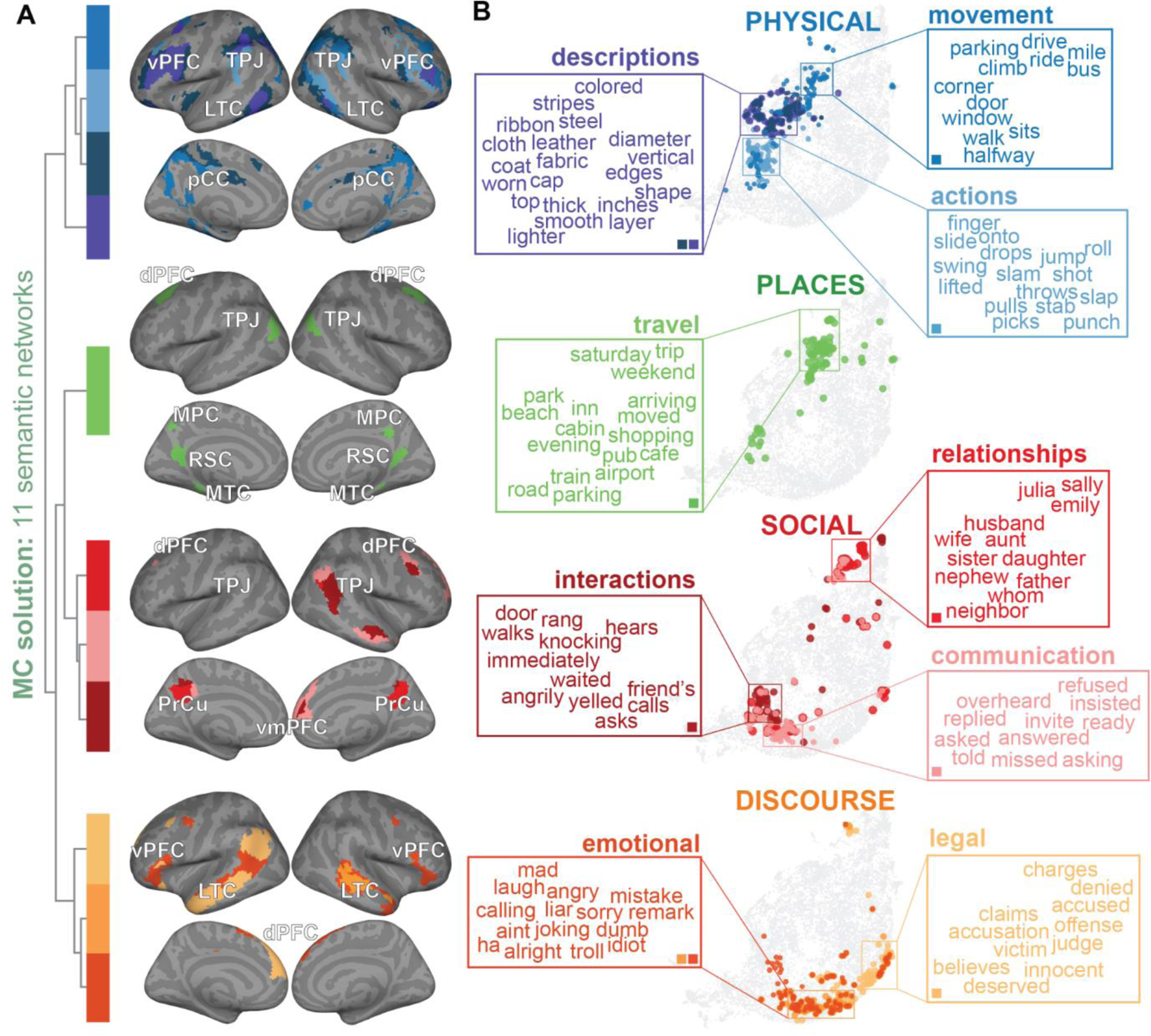
Model connectivity recover networks with rich and precise functional definitions. MC recovers networks based on the encoding model weights that define the functional tuning of each vertex. We can therefore directly interpret each network in terms of the average functional tuning across vertices within the network. (**A**) The 11 semantic networks recovered by MC are split into four groups (as in Figure 3A), and are visualized on the inflated template surface. The 12-th network is not reliably predicted by the semantic encoding models, so it is excluded from interpretation. (**B**) To define the semantic tuning of each network, we identified the 100 words that were predicted to elicit the largest BOLD response in each network. These 100 words are plotted here in a 2D semantic space generated by UMAP. In this 2D semantic space, each point represents one of the words spoken in the stories, and neighboring points are semantically similar words. The size of each point corresponds to the magnitude of the predicted BOLD response to the word, and the color of each point corresponds to its network assignments. The insets show a representative sampling of the 100 words (e.g., colored, stripes, steel) plotted as a word cloud for each network. The colored squares inside each inset indicate the networks that correspond to the depicted semantic tuning. The semantic category represented by each network was interpreted by manually inspecting these 100 words (e.g., descriptions), and is written above each word cloud. From this approach, we determined that nine distinct categories of physical-, place-, social-, and discourse-related semantic concepts are represented by the 11 semantic networks. This analysis demonstrates that the encoding model weights provide each network with a rich functional description that describes the functional contribution of the network during the task.

To interpret the precise category of semantic concepts represented by each semantic network, we manually inspected the 100 words that were predicted to elicit the largest BOLD response in the network. This process revealed that the four networks colored in shades of blue represent physical-related categories of semantic concepts: two networks represent physical descriptions (e.g., colored, inches), one network represents physical movement (e.g., climb, drive), and one network represents physical actions (e.g., swing, punch). The single network colored in green represents a place-related category: travel (e.g., trip, beach). The three networks colored in shades of red represent social-related categories: one network represents social interactions (e.g., knocking, waited), one network represents social relationships (e.g., wife, father), and one network represents communication (e.g., replied, asking). The three networks colored in shades of orange represent discourse-related categories: two networks represent emotional discourse (e.g., laugh, troll) and one network represents legal discourse (e.g., claims, judge). While the interpretation of the precise function of each network is necessarily subjective, the semantic encoding model weights provide rich descriptions of the functional selectivity of each network. Thus, MC recovers networks with highly interpretable functional descriptions.

Because each of the networks have a rich profile of semantic tuning, these semantic networks also provide insight into the functional tuning of the canonical resting state networks during natural language comprehension (see **Figure 5B** for the most spatially similar MC and resting state networks). The two MC networks that were most similar to the canonical attention networks recovered by Yeo et al.^32^ represent physical actions and physical descriptions. The two MC networks that were most similar to the canonical frontoparietal control networks represent physical movement and physical descriptions. The single MC network that was most similar to the temporoparietal network represents travel. Finally, the six MC networks that subdivide the DMN represent communication, social relationships, social interactions, legal discourse, and emotional discourse. These analyses suggest that during natural language comprehension, the canonical attention, frontoparietal control, and temporoparietal networks represent concrete, physically-grounded categories of semantic concepts, whereas the canonical DMN represents abstract, social-related categories of semantic concepts. Thus, by leveraging the advantages provided by MC, we can determine the functional role of canonical resting state networks during a task.

### Networks recovered by MC have similar functional tuning across individuals

Because the 12-network MC solution optimized cross-participants prediction accuracy, this group solution reflects patterns of functional coupling that are shared across participants. Given the anatomical and functional differences across individuals^58^, these 12 MC networks might not reflect the functional organization of language comprehension systems in individual participants. To test how closely the group solution captures functional tuning in individuals, voxelwise semantic encoding models were fit to functional data from each participant in their native anatomical space, and were then spatially mapped to each participant’s cortical surface. The group 12-network MC solution was also mapped to each participant’s cortical surface (**Figure 7A**), and the participant’s model weights were averaged across vertices within each spatially mapped network. This approach allowed us to recover and compare the functional tuning of the group MC networks in individual participants.

**Figure 7.**
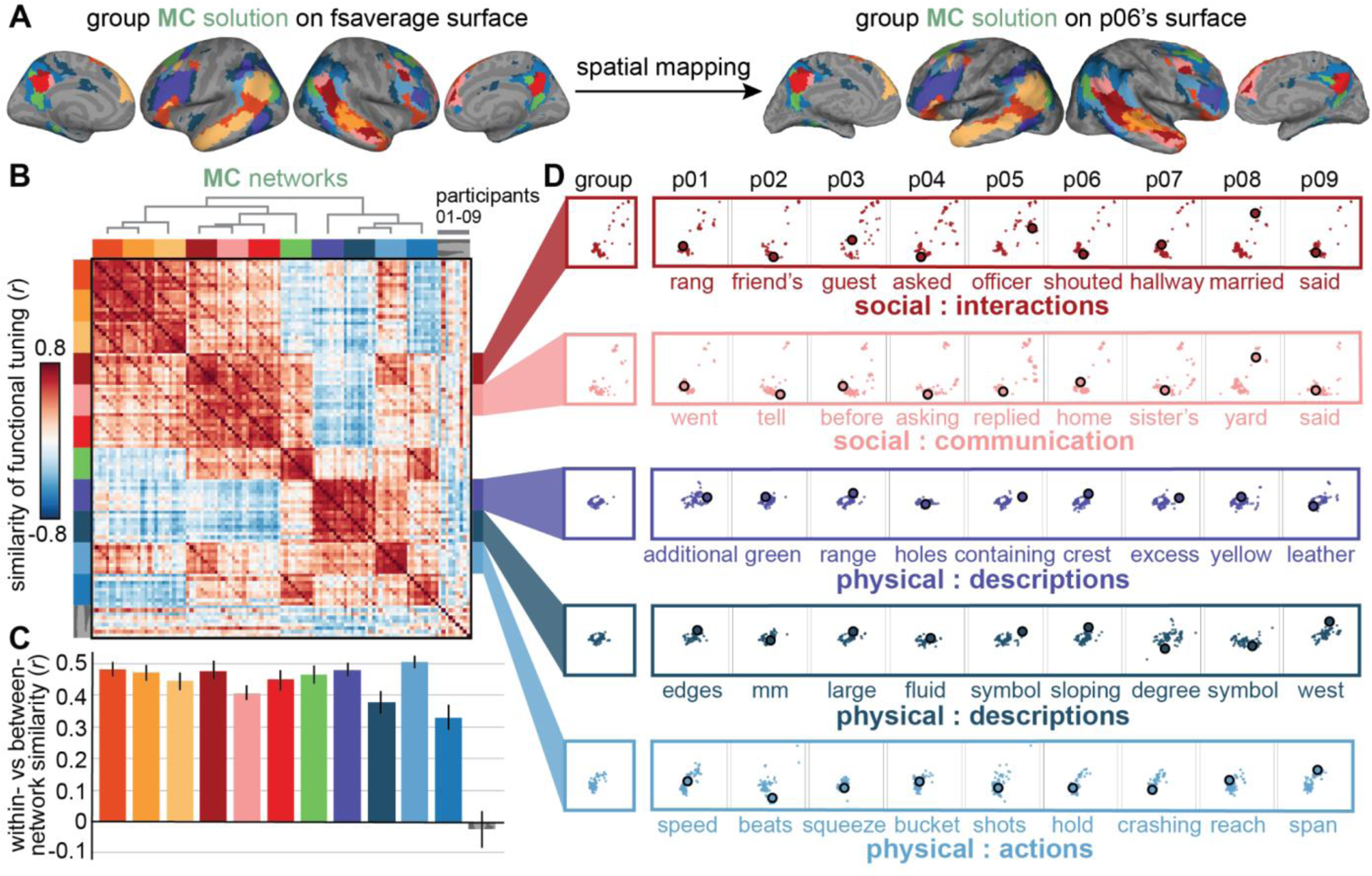
Spatially mapping the group MC solution to each participant’s cortical surface reveals that the group networks have consistent functional tuning across participants. Mapping individual participants’ data to a template surface and averaging across participants recovers a 12-network MC solution that reflects the functional organization of semantic processing systems at the group level. To evaluate how well the group solution reflects functional organization within each participant, we mapped the group solution to each participants’ cortical surface to assess functional tuning within each individual participant. (**A**) The group MC solution was mapped from the fsaverage template surface to each participant’s cortical surface. The resulting spatially mapped MC solution is shown for participant 06 on their inflated cortical surface. This spatial mapping preserves the spatial organization but not necessarily the functional tuning of each group MC network in each individual participant. (**B**) To quantify the consistency of semantic tuning across individuals, the network-average semantic model weights were first correlated across participants and then aggregated in a pairwise correlation matrix. The rows and columns of the matrix are grouped by network and ordered by participant. The block structure along the main diagonal suggests that, across participants, semantic weights are highly similar within a network. (**C**) The bar graph shows the difference between the cross-participant consistency within a network and the cross-participant consistency between networks. The color of each bar corresponds to the network assignment, and the error bars indicate the bootstrapped 95% confidence interval. Across participants, the semantic tuning within the 11 semantic networks is significantly more similar within the network than between networks (*p* < 0.001, permutation test). By contrast, the semantic tuning within the “noise” network is as consistent within the network than between networks (*p* = 0.70). (**D**) To visualize the consistency of functional tuning within networks and across participants, the semantic coverage of five networks is plotted separately for each participant (denoted as p01-p09). Each participant’s network-average model weights were used to identify the words that were predicted to elicit the strongest BOLD response in the network. The top 100 words are plotted as points in a 2D semantic embedding space (as in Figure 6B). For each network and each participant, one word that is representative of the semantic tuning of the network is highlighted. Within each network, the distribution of words in the semantic embedding space largely overlaps across participants. Thus, the group MC networks represent similar categories of semantic concepts across individuals.

To quantify the cross-participant consistency within each network, each participant’s network-average model weights were correlated across networks and across participants (**Figure 7B**). The blocks of high correlation along the main diagonal of the pairwise correlation matrix suggest that the model weights are more similar within the network than between networks. **Figure 7C** shows the difference between the within-network correlations and the between-network correlations for each network. For each of the 11 semantic networks, the cross-participant semantic tuning within the network is significantly more consistent than the cross-participant semantic tuning between networks (difference = 0.44 ± 0.05, mean ± s.d. across networks; *p* < 0.001 for each semantic network, permutation test). By contrast, the cross-participant semantic tuning within the noise network is not significantly different from the cross-participant semantic tuning between the noise network and the 11 semantic networks (difference = −0.03; *p* = 0.70). These analyses indicate that the networks that are well-predicted by the encoding models also have consistent functional tuning across participants.

To visualize the cross-participant consistency of functional tuning within each network, the range of words represented by five example networks were plotted in the two-dimensional semantic space, both in the group (labeled in **Figure 7D** as group) and in each individual participant (labeled in **Figure 7D** as p01–p09; see **Supplemental Figure 5** for the functional tuning of each network in each participant). For each of the five networks, the 100 words that were predicted to elicit the largest BOLD response are plotted in the same 2D semantic space as in **Figure 6B**. For the group, the top 100 words were recovered using model weights averaged across the group of participants. For each individual, the top 100 words were recovered using model weights estimated in the participant’s native anatomical space. Across participants, a similar set of words are predicted to elicit the largest change in BOLD responses within each network (proportion of the top 100 words that are shared between the group and participant-level models = 0.52 ± 0.18, mean ± s.d. across participants). Thus, the 11 semantic MC networks recovered from the group largely capture the functional organization in individual participants.

When examining the functional tuning profiles of the networks in individual participants, we noticed that the pairs of networks with very similar functional tuning at the group level show differences in functional tuning across individuals. For instance, at the group level, the functional tuning of the dark purple and dark blue networks is indistinguishable (see group column in **Figure 7C**). However, examining the functional tuning of each network in each individual participant reveals that the functional tuning within the dark blue network is less consistent across individuals (within-network *r* = 0.55 ± 0.11, mean ± s.d. across pairwise comparisons) than the functional tuning within the dark purple network (*r* = 0.64 ± 0.06), a statistically significant difference (difference = 0.09, *p* < 0.001, permutation test). This suggests that the group MC networks reflect both differences in the functional tuning and differences in the cross-participant consistency of functional tuning.

We also observed that pairs of networks that share a spatial border on the cortical surface often have similar functional tuning. For example, functional tuning is correlated between the network that represents social interactions colored in dark red and the network that represents physical actions colored light blue (*r* = 0.44 ± 0.11, mean ± s.d. across pairwise comparisons), which share spatial borders on the cortical surface (see **Figure 3A**). This suggests that some of the similarity in functional tuning between pairs of networks might be caused by individual differences in the cortical distribution of these networks. When two networks neighbor each other on the cortical surface, individual differences in the cortical extent of the networks blur the functional tuning within each network, thereby increasing the similarity of functional tuning between neighboring networks.

Overall, these analyses suggest that the functional tuning within the group MC networks are largely consistent across individuals. However, spatially mapping the group MC solution to each participant’s cortical surface does not appear to adequately capture differences in anatomical or functional organization that exist across individuals.

### Networks recovered by MC have similar cortical distributions across individuals

In the previous section, the group MC solution was recovered in individual participants by spatially mapping the solution from the template cortical surface to each participant’s cortical surface (generating *spatially mapped MC solutions*). While this process preserves the spatial organization of the functional networks recovered from the group, it cannot account for individual differences in anatomical or functional organization. To recover these differences, we implemented another approach for recovering the MC networks in each participant that better preserves anatomical and functional differences across individuals (**Figure 8A**). First, the group-average model weights were averaged across vertices within each network of the group solution. Then, the correlation was computed between the network-average model weights of the group solution and the vertex-wise model weights of each participant. Lastly, each vertex in each participant was assigned to the MC network with the most similar functional tuning, recovering the group MC networks in each individual participant (generating *functionally matched MC solutions*). Differences in the spatially mapped and functionally matched networks reveal which cortical regions contain the most variable functional organization across individuals.

**Figure 8.**
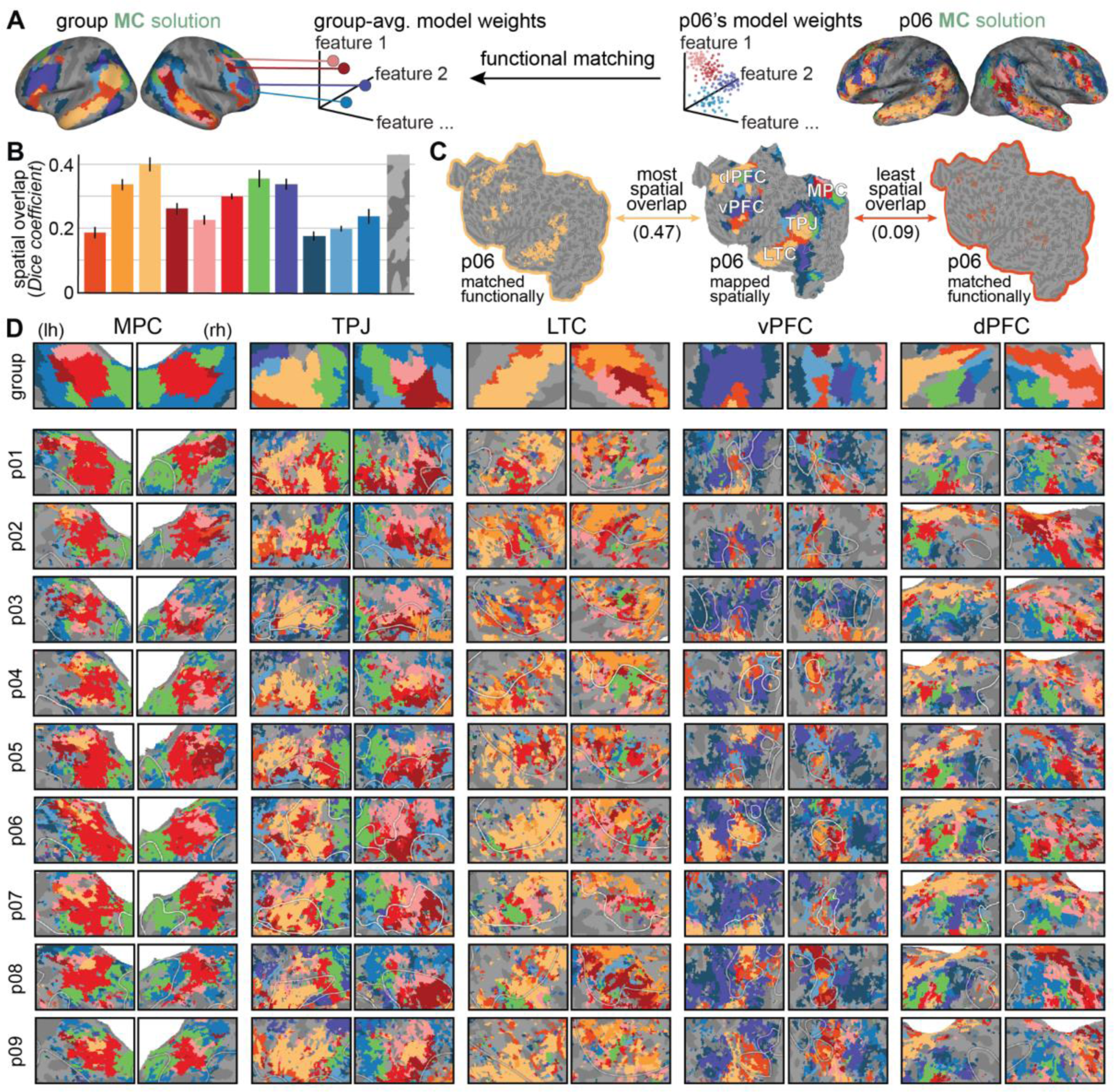
Functionally matching each vertex of each participant’s cortical surface to one of the group MC networks recovers individual differences in functional organization across participants. Spatially mapping the group MC solution to each participant’s cortical surface ignores individual differences in anatomical and functional organization. To better account for such individual differences, we used the group network-average functional tuning to recover the group MC networks in individual participants. (**A**) To recover the group MC networks in each participant, the group-average model weights were first averaged across vertices within each MC network on the template surface. The model weights of each vertex of each participant were then compared to the group network-average model weights, and each vertex was assigned to the MC network with the most similar functional tuning. The resulting functionally matched MC solution is shown for participant 06 on their inflated cortical surface. This functional matching preserves the functional tuning but not necessarily the cortical distribution of each group MC network in individual participants (see **Supplemental** Figure 6 for the functionally matched MC solution in each participant). (**B**) To assess how variable each group MC network is across participants, the Dice coefficient was used to quantify the spatial overlap between the spatially mapped MC networks and the corresponding functionally matched MC networks. The color of each bar indicates the network assignment, and the error bars indicate the s.e.m. across participants. Variability in the degree of spatial overlap between the spatially mapped and functionally matched MC networks indicates that some group networks (e.g., the networks colored in light orange and green) have a more consistent cortical distribution across individuals, whereas other networks (e.g., the networks colored in dark orange and dark blue) have a more variable cortical distribution across individuals. (**C**) To visualize the range of spatial overlap between the functionally matched networks (left and right) and the spatially mapped networks (center), the functionally matched network that overlaps most with the corresponding spatially mapped network (left, outlined in light orange) and the functionally matched network that overlaps least with the corresponding spatially mapped network (right, outlined in dark orange) are shown for the left hemisphere of participant 06’s flattened cortical surface. The corresponding Dice coefficients are indicated in parentheses. While the sparsity differs between the network with the most spatial overlap and the network with the least spatial overlap, both networks span a similar range of cortical areas in the functionally matched and spatially mapped solutions. This suggests that even group networks with relatively low degree of spatial overlap are recovered in individual participants. (**D**) The MC networks within the left and right hemispheres (labeled lh and rh) of five distinct cortical regions (labeled MPC, TPJ, LTC, vPFC, and dPFC) are shown on the flattened cortical surface. The group MC solution is shown on the template surface at the top (labeled group), and the functionally matched MC solutions of all nine participants are shown on each participant’s cortical surface below (labeled p01-p09). While the cortical distribution of the group- and participant-level MC networks are largely consistent across these cortical regions, the participant-level networks capture individual differences in functional organization that are not captured by the spatially mapped networks. Thus, functionally matching networks allows us to recover individual differences in the cortical distribution of the group-level functional networks.

To quantify the variability of each group MC network across individuals, the Dice coefficient was computed between the spatially mapped and functionally matched MC networks (**Figure 8B**). A high Dice coefficient indicates that the cortical distribution of the group-level MC network is consistent across individuals, whereas a low Dice coefficient indicates that the cortical distribution of the group-level MC network varies across individuals. Across participants, the most consistent networks include the group MC network that represents semantic concepts related to legal discourse (colored in light orange; *D* = 0.40 ± 0.02, mean ± s.e.m. across participants), and the group MC network that represents concepts related to places (colored in green; *D* = 0.35 ± 0.03). By contrast, the most variable networks include one of the group MC networks that represents emotional discourse (colored in dark orange; *D* = 0.19 ± 0.02), and one of the group MC networks that represents physical descriptions (colored in dark blue; *D* = 0.17 ± 0.02). These results suggest that some group MC networks have a more consistent cortical distribution across individual participants than other group MC networks. To visualize the range of spatial variability observed across group MC networks, **Figure 8C** shows the network with the most spatial overlap in participant 06 (colored in light orange; *D* = 0.47) and the network with the least spatial overlap in participant 06 (colored in dark orange; *D* = 0.09). While the network with the least spatial overlap is sparser than the network with the most spatial overlap, both networks have a similar cortical distribution between the spatially mapped and functionally matched solutions. This suggests that even group MC networks with relatively minimal spatial overlap between the spatially mapped and functionally matched solutions can be recovered in individual participants.

The results of these analyses are largely consistent with our findings from the previous section, which suggest that the group MC networks reflect both differences in functional tuning and differences in the consistency of functional organization across participants. However, because this functional matching approach does not constrain the cortical distribution of the participant-level networks, it also reveals the cortical regions where functional organization varies the most across individuals. To qualitatively assess which cortical regions have a consistent functional organization across participants, we compared the functional organization determined by the group MC solution and the participant-level MC solutions across five distinct cortical regions (**Figure 8D**; see **Supplemental Figure 6** for the participant-level MC solutions recovered in each individual participant). The group MC solution appears to capture much of the broad functional organization that is preserved across participants, a result consistent with that shown in **Figure 7**. However, this functional matching approach also reveals that the functional organization within the MPC and TPJ appears to be more consistent across participants, whereas the functional organization within the ventral PFC (vPFC) and dorsal PFC (dPFC) appears to be more variable across participants. Thus, by using two distinct approaches to recover the group MC networks in individual participants, we can recover both the broad functional organization that generalizes across participants, and the variations in functional organization that are specific to each individual.

## Discussion

Model connectivity (MC) is a powerful new method for recovering functional brain networks that overcomes the flaws in functional connectivity (FC). The flaws in FC stem from two fundamental characteristics of the method. First, FC assumes that all temporally correlated brain activity reflects true functional coupling. We showed that applying FC to an fMRI dataset collected during a natural language comprehension task produces functional networks that are confounded by differences in SNR across the brain. Second, FC does not provide any means to define the functional information reflected in the measured BOLD activity. We showed that the networks recovered by FC can only be assigned a coarse functional description that is not specific to the stimulus or task. To overcome these flaws, MC leverages the power of the Voxelwise Modeling framework, recovering networks of brain regions with similar encoding model weights. Applying MC to the same narrative auditory dataset as FC produces functional networks that capture stimulus- or task-specific patterns of functional coupling that are shared across participants. Additionally, the associated encoding model weights provide each network with a rich functional description in terms of the stimulus- or task-related features. Thus, the networks recovered by MC define the functional organization that underlies complex cognitive functions and behaviors.

### FC is confounded by spatial variability in SNR

It is well known from studies of temporal SNR that the relative contribution of signal and noise varies systematically across the brain^13, 59^. Because FC is based on temporally correlated fluctuations in BOLD activity, spatial differences in SNR inevitably influence the patterns of functional coupling measured by FC. The majority of FC studies use complex preprocessing pipelines in an attempt to reduce temporal noise in the measured BOLD activity^5, 10, 60–62^, but these approaches do not account for the spatial differences in SNR that are caused by anatomy, such as head and brain geometry, vasculature, and differences in magnetic susceptibility. In an attempt to account for broad spatial differences in SNR, some FC studies exclude brain regions with a temporal SNR that is below a certain threshold from their analyses^32, 63^. However, even subtle differences in tSNR contaminate the networks recovered by FC (see **Figure 4** and **Supplemental Figure 4**), indicating that a simple tSNR threshold cannot fully correct for these confounds.

The problem of noise contaminations is especially pronounced for studies that compare FC across participant groups^64–69^ or across participants^20, 63, 70–72^, as noise contributions vary substantially across individuals. For instance, previous studies have noted that differences in the amount of motion between participant groups can be incorrectly interpreted as differences in functional coupling^60, 61^. Furthermore, patterns of spatial variability in SNR are unique to individual participants (see **Supplemental Figure 4** for spatial patterns of tSNR in each participant), and are rarely accounted for in FC studies (see Baker et al.^68^ for an approach that accounts for differences in noise levels across participant groups, but not for differences in spatial patterns of SNR). FC studies are therefore prone to misinterpreting differences in SNR between participant groups or between participants as meaningful differences in functional coupling.

MC overcomes this flaw in FC by ignoring fluctuations in BOLD activity that do not reflect reliable functional activity. The encoding model weights used by MC quantify only the stimulus- or task-related information that is represented in the BOLD activity. Therefore, because MC is based on encoding model weights, it only reflects fluctuations in BOLD activity that are related to true stimulus- or task-related functional activity.

### FC is not directly interpretable

The overarching goal of task-based FC studies is to assess how patterns of functional coupling support specific cognitive functions or behaviors. However, because FC provides no means to identify the specific aspects of the stimuli or tasks that result in the measured patterns of functional connectivity, task-based FC studies rely on additional datasets to interpret the functions of the recovered networks. Specifically, most task-based FC studies reference the canonical set of networks recovered from resting state functional data that are posited to reflect intrinsic patterns of functional coupling^32, 33^. Task-based FC studies treat these resting state networks as the baseline functional organization of the brain, and assume that the task-based networks are reconfigurations of these resting state networks^73^. However, variations in the cortical distribution of the networks may be caused by factors that are not related to meaningful differences in functional coupling between the task and rest. For example, anatomical or functional differences between the participants in each dataset^56, 57, 70–72, 74, 75^, and transient differences in the participants’ attentional or arousal state^76–78^ have both been reported to influence the patterns of functional coupling measured by FC. Furthermore, because attention changes representations across the brain^79^, the precise functional role of each of the canonical resting state networks may change according to the task. The functional domain assigned to each resting state network may therefore be misleading^34^. Thus, FC studies cannot assign functional significance to differences observed in FC between task and rest.

MC overcomes this flaw in FC by using the same dataset to recover and interpret the functional networks. These networks explicitly represent the stimulus- or task-related information that underlies a specific component of behavior. In the present study, the semantic encoding model weights that were used to recover distinct functional networks were also used to identify the precise category of semantic concepts represented by each network. Therefore, the resulting semantic networks reveal both the functional organization of semantic processing systems in the brain, and the functional tuning of each of these systems.

Because we were able to directly interpret the function of each MC network, applying MC to the narrative auditory dataset provides insight into the functional role of canonical resting state networks during natural language comprehension. Specifically, half of the networks recovered by MC were most similar in cortical distribution to the canonical DMN recovered by Yeo et al.^32^. Examining the functional tuning of these six semantic networks revealed that they represent social- and discourse-related categories of semantic concepts. This finding suggests that the DMN represents abstract, social semantic concepts during language comprehension. Previous studies have implicated the DMN in a variety of complex cognitive processes, including introspection^80, 81^, memory^82, 83^, social reasoning^84, 85^, internal speech^86, 87^, and processing conceptual semantics^40, 88–90^. Therefore, the overarching role of the DMN may be representing abstract, social-related concepts, regardless of whether the concepts are internally generated or externally perceived.

### FC is insensitive to task-based differences in functional coupling

Given that the networks recovered by FC are confounded by systematic differences in SNR and are limited in their functional interpretability, we argue that FC is not well suited for examining the differences in functional coupling across tasks. This view is largely supported by previous FC studies that compare FC between task and resting state. Studies that only looked at group-level effects reported a minimal influence of task state on the measured FC between brain areas^73, 91^. Furthermore, while studies that compared FC across individuals and tasks noted a larger effect of task state^63, 71, 92, 93^, the magnitude of the effect of individual differences far outweighed the magnitude of the effect of task-based differences^63, 94^. These studies argue that stability across task and resting states is a benefit of FC, indicating that FC captures the state-invariant functional architecture of the brain. However, we argue that this stability may also reflect a limitation of using FC for task-based studies, indicating that it captures reliable spatial patterns of BOLD SNR, which are not related to functional processes, rather than the stimulus- or task-specific information that is represented in the BOLD activity.

MC overcomes this limitation of FC by evaluating functional coupling in terms of stimulus- or task-related features related to the experimental paradigm. This makes MC an ideal method for comparing how the whole-brain patterns of functional coupling support the perceptual, cognitive, and motor aspects of complex behaviors. For example, different feature spaces capture unique stimulus- or task-related information^37, 44^. Therefore, applying MC separately to the high-level semantic model weights and to the low-level auditory model weights would recover distinct sets of networks, depicting the functional organization of the high-level cognitive and the low-level perceptual representations necessary for natural language comprehension. Furthermore, MC allows us to examine how these whole-brain patterns of functional coupling change according to experimental conditions. For example, because the set of brain areas that are involved in processing semantic information differs between auditory and visual modalities^95^, applying MC to the semantic model weights estimated during story listening and to the semantic model weights estimated during movie watching could recover distinct sets of networks. Additionally, because top-down selective attention widely affects functional tuning across the cortical surface^79^, applying MC to model weights estimated under different attentional states could reveal how attention affects the whole-brain functional organization. Thus, by using MC to compare both the cortical distribution and the functional tuning of networks across feature spaces, perceptual modalities, and attentional states, we can discover how changes in whole-brain patterns of functional coupling support the specific perceptual, cognitive, or motor components of an experimental task.

### MC provides a complete framework for recovering functional networks from encoding model weights

The present study is not the first to group brain areas based on encoding model weights. The original publication of this dataset used a Bayesian algorithm called PrAGMATiC to recover a set of functionally homogeneous brain areas that tiled the cortical surface^40^ (see **Supplemental Figure 7** for a comparison in prediction accuracy between the group MC solution recovered in the present work and the group PrAGMATiC solution). However, PrAGMATiC is complex and computationally costly. Several other studies took an approach similar to that used here, clustering voxels based on encoding model weights^96–98^. However, those studies did not develop a systematic approach for determining the number of clusters that best generalize to held-out participants. Therefore, the clusters identified in those studies may miss functional subdivisions that exist in the data. By contrast, MC provides a simple, robust framework for recovering the full set of networks that most accurately reflect the functional organization of the brain during an experimental task.

## Conclusion

Model connectivity is a powerful, data-driven method for recovering functional networks that overcomes the limitations of functional connectivity. Because FC is based solely on temporally correlated BOLD activity, FC does not distinguish between signal and noise in the BOLD activity. Consequently, FC recovers networks that are confounded by spatial variability in SNR, and that are difficult to interpret. These flaws make FC poorly suited for studying differences in functional coupling across tasks or experimental conditions. In contrast, MC is based on an explicit model of the repeatable functional signal in the BOLD activity. Thus, MC recovers networks that reflect the underlying functional organization, and that have rich functional descriptions that are directly related to the stimulus or task. These qualities make MC an ideal method for examining how the cortical distribution and the functional tuning of whole-brain functional networks support complex tasks and behaviors. To encourage the adoption of MC, the code implementing the full pipeline is openly available as a Python package [link will be provided here upon publication].

## Acknowledgments

We thank Catherine Chen for her suggestions for recovering the group networks in individual participants. We thank the members of the Gallant Lab for aiding in data collection and flattening cortical surfaces. This work was supported by grants from the National Science Foundation (NSF; IIS1208203), the National Eye Institute (EY019684 and EY031455), the National Institute on Aging (AG029577), and the Office of Naval Research (N00014-22-1-2217).

## Author contributions

Conceptualization, M.V.d.O.C. and E.X.M.; Methodology, E.X.M., M.V.d.O.C., and T.D.l.T; Software, E.X.M. and T.D.l.T; Investigation, E.X.M.; Visualization, E.X.M.; Writing – Original Draft, E.X.M.; Writing – Reviewing & Editing, E.X.M., M.V.d.O.C., T.D.l.T, and J.L.G.; Data Curation, M.V.d.O.C.; Supervision, M.V.d.O.C. and J.L.G.; Funding Acquisition, J.L.G.

## Methods

### Participants

This study was approved by the Committee for Protection of Human participants at the University of California, Berkeley. All participants gave written informed consent. Functional data were collected from 11 participants (3 females) between the ages of 24 and 33. Data from two participants were held-out for validation. Data from 6 of the 11 participants used in this work were originally collected for a previously published project^40^. Since the original publication, data were collected from an additional 5 participants, using the same experimental paradigm as used in Huth et al.

All voxel- and vertex-wise encoding models were estimated and validated independently for each participant using separate data sets. Data were averaged across 9 of the 11 participants to recover functional networks. Across the 9 participants, a leave-one-participant-out cross-validation scheme was used to evaluate whether the group networks accurately predict brain activity from each participant. The remaining two participants were used to evaluate how well the recovered networks predict brain activity from completely held-out participants (**Supplemental Figure 7**).

### Experimental procedure

Participants listened to eleven 10- to 15-min stories taken from *The Moth Radio Hour*. In each story, a speaker tells an autobiographical story in front of a live audience. The eleven selected stories cover a wide range of topics and are highly engaging. Each story was played during a separate fMRI scan. The length of each scan was tailored to the story and included 10 s of silence both before and after the story.

The eleven stories from *The Moth Radio Hour* were split into a model training set and a model test set. The training set consisted of ten 10- to 15-min stories, and contained a total of 125-min of data for each participant. The test set consisted of one 10-min story that was presented to each participant two separate times. To reduce noise in the test dataset, the BOLD responses to the two repetitions of the test story were averaged for each participant. The final test set contained a total of 10-min of data for each participant.

### MRI data collection

MRI data were collected on a 3T Siemens Tim Trio scanner at the University of California, Berkeley Brain Imaging Center using a 32-channel Siemens head coil. Functional scans were collected using gradient echo EPI with repetition time (TR) = 2.0045 s, echo time (TE) = 31 ms, flip angle = 70°, voxel size = 2.24 × 2.24 × 4.1 mm (slice thickness = 3.5 mm with 18% slice gap), matrix size = 100 × 100 and field of view = 224 mm × 224 mm. Thirty axial slices covered the entire cortex and were acquired in ascending interleaved order. A custom-modified bipolar water excitation radiofrequency pulse was used to avoid signal from fat. For most subjects (9 out of 11), a dual-echo gradient echo fieldmap was acquired before every functional scan. Anatomical scans were collected using a T1-weighted multi-echo MP-RAGE sequence with the parameters recommended by FreeSurfer.

### MRI data preprocessing

#### Preprocessing steps

Each functional run was motion-corrected, distortion-corrected, and aligned to the participant’s T1 scan. Motion correction transformations were computed with the FMRIB Linear Image Registration Tool (FLIRT) from FSL^99, 100^. Distortion correction transformations were computed with FUGUE from FSL^101^. Alignment transformations were computed with bbregister from FreeSurfer^102^. Finally, all transformations were combined and applied in a single transformation step using ANTs^103^. The resulting preprocessed data were then temporally detrended and denoised using CompCor^104^ and RETROICOR^105^. Note that this preprocessing pipeline differs from the pipeline used in our previous publications. This new pipeline adopts several improvements first developed and validated by fMRIPrep^106^, such as using bbregister and ANTs to align functional images to each participant’s native space.

#### Projection to fsaverage5

FreeSurfer’s *mri_vol2surf* was used to project the pre-processed functional volumes to the fsaverage5 surface, which results in a minimal amount of smoothing (∼3 mm full-width half-max kernel). The fsaverage5 template surface consists of 10,242 vertices per hemisphere that are spaced approximately 3 mm apart (20,484 vertices total). In fsaverage5 space, functional data within each run were z-scored over time.

#### Visualization

Functional data were visualized on the flattened and inflated cortical surface using pycortex, a Python package developed by our lab^107^. All analyses were performed on data projected to the fsaverage5 surface. The results of the analyses were projected to the high-resolution fsaverage surface for visualization only (327,684 vertices total).

### Encoding model estimation

Encoding models use features extracted from the experimental stimulus or task to predict brain activity in each participant. In the present work, high-level semantic features and low-level auditory features were extracted from the narrative auditory stories. Regularized linear regression was used to predict brain activity from the semantic and auditory features. A separate encoding model was estimated for each vertex of the template cortical surface, so we refer to these models as *vertex-wise* encoding models. Separate training and test datasets were used to estimate and evaluate vertex-wise encoding models for each participant.

#### Semantic features

Semantic features were derived from co-occurrence statistics between the set of 10,470 words spoken across the ten stories and a set of 985 common English words^40^. Co-occurrence rates were calculated on a large corpus of text that included 13 Moth stories, 604 popular books, 2,405,569 Wikipedia pages, and 36,333,459 user comments scraped from reddit.com. The 10,470 spoken words appeared a total of 1,548,774,960 times in this corpus. The word co-occurrence matrix, *M*, contains 985 rows and 10,470 columns. Each element of *M* represents the proportion of times the spoken word occurs within 15 words of the common word. Each column and row were z-scored to account for differences in how often each word appeared in the corpus. Thus each column of *M* represents a 985-dimensional semantic embedding for each of the 10,470 spoken words.

The features used for model estimation were then constructed from the ten training stories for each word-time pair (*w*, *t*). For each word in each story, the corresponding column of *M* representing the spoken word’s 985-dimensional semantic embedding was appended. These vectors were then resampled at times corresponding to the fMRI acquisitions using a 3-lobe Lanczos filter with the cut-off frequency set to the Nyquist frequency of the fMRI acquisition (0.249 Hz).

#### Low-level auditory features

Low-level auditory features were included to account for responses to low-level properties of the stories that could contaminate the semantic model weights. The 41 low-level auditory features were word rate (one feature), phoneme rate (one feature) and phonemes (39 features).

#### Feature adjustments

Before fitting, each stimulus feature within each story run was demeaned over time. To account for the hemodynamic delay, a separate linear temporal filter with four delays (1, 2, 3 and 4 time points) was fit for each feature. This was accomplished by concatenating feature vectors that had been delayed by 1, 2, 3 and 4 time points (2 s, 4 s, 6 s and 8 s). For instance, in the concatenated “word rate” feature space, one feature represents the word rate 2 s earlier, another 4 s earlier, and so on. Taking the dot product of this concatenated feature space with a set of linear weights is functionally equivalent to convolving the original stimulus vectors with linear temporal kernels that have non-zero entries for 1, 2, 3 and 4 time point delays.

#### Model estimation

Models were fit to functional data independently for each participant in each vertex of the fsaverage5 surface. A modified version of ridge regression called banded ridge regression was used to model the functional data^43, 44^. Two separate regularization hyperparameters were optimized for each vertex: one for the semantic features and one for the low-level features. Separately optimizing these hyperparameters for each vertex allows the model to weigh the two sets of features independently across the brain.

The optimal regularization hyperparameters were determined via cross-validation. The model training set, which consisted of BOLD responses to ten auditory stories, was split into a smaller training and validation set. The smaller training set consisted of nine stories, and the validation set consisted of one story (leave-one-run-out cross-validation scheme). Then, multiple banded ridge regression models were fit with 20 logarithmically spaced regularization hyperparameters ranging from 10^1^ to 10^20^. The prediction accuracy of each of these models was estimated on the validation set. This process was repeated by leaving out each of the ten model training stories. The resulting prediction accuracies were then averaged across left-out stories, and the hyperparameters with the highest prediction accuracy were selected separately for each vertex.

All model fitting was performed using himalaya, a Python package developed by our lab to efficiently fit ridge and banded ridge regression models^44^.

#### Model evaluation

The prediction accuracies of the final vertex-wise encoding models were evaluated for each participant using the model test dataset. For each vertex, model prediction accuracy was evaluated as the Pearson’s correlation coefficient (*r*) between the predicted BOLD responses and the recorded BOLD responses.

### Recovering functional networks

Model connectivity and functional connectivity were used to recover networks at the group level. In both approaches, connectivity was assessed between all 20,484 vertices of the fsaverage5 surface. For MC, model weights were estimated independently in each vertex for each participant, and averaged across participants (we refer to these averaged weights as *group-average model weights*). Connectivity between vertices was then evaluated as the Euclidean distance between group-average model weights. For FC, BOLD time-series in each vertex were averaged across participants (we refer to these averaged time-series as *group-average BOLD responses*). Connectivity between vertices was then evaluated as the correlation distance (1 - correlation) between group-average BOLD responses. The pairwise distances between all vertices were stored in two vertex-wise 20,484 x 20,484 distance matrices, one for MC and one for FC. To group vertices into networks, hierarchical clustering with Ward’s linkage was applied to each distance matrix^53, 108, 109^. To limit overfitting, networks recovered by MC were based on model weights estimated using the training dataset (125-min, 3737 TRs).Similarly, networks recovered by FC were based on BOLD time-series from the training dataset. For both MC and FC, networks were evaluated using the test dataset (10-min, 291 TRs).

#### Vertex-wise clustering

FC studies employ a wide variety of methods to recover functional networks, including seed-, graph-theory-, and clustering-based approaches^110, 111^. In the present work, we used a vertex-wise clustering approach to recover MC and FC networks, as clustering enforces the fewest assumptions about the spatial organization of the recovered networks. Seed-based approaches rely on defining a set of seed regions of interest^6^, and graph-theory-based approaches rely on defining a subset of functionally homogeneous parcels or nodes^112^. These seed regions and sets of spatially contiguous parcels are assumed to be stable across task states, and to contain similar time-courses of BOLD activity^49–51, 113^. However in many cases, these assumptions are too strong. For any given parcel, both the cortical extent and the functional homogeneity within the parcel may differ across groups, individuals, or task states^93, 114^. Using a parcellation recovered from a separate dataset may therefore bias the results. By contrast, clustering-based approaches do not rely on defining a smaller subset of regions of interest or functionally homogeneous parcels. Thus, by clustering all possible vertices of the cortical surface, the recovered networks reflect the current data rather than prior spatial constraints.

#### Selecting the distance metric

Two different distance metrics were used to recover functional networks under MC and FC. MC used Euclidean distances of encoding model weights in a feature space, and FC used correlation distances of BOLD activity over time. These two distance metrics reflect different assumptions about how MC and FC define functional coupling.

Under MC, functional coupling is assumed to be determined by the similarity of functional tuning between brain areas. Functional tuning is quantified by the encoding model weights, which define a vector in the feature space. This vector can be decomposed into a magnitude and a direction in the feature space, both of which contribute to the functional tuning profile. Because the Euclidean distance preserves the contribution of both magnitude and direction, MC uses the Euclidean distance to quantify functional coupling.

Under FC, functional coupling is assumed to be determined only by temporal co-fluctuations in BOLD activity between brain areas. In FC, the magnitude of the BOLD activity is not considered to carry any meaningful information about functional coupling. Because the correlation distance measures the degree to which BOLD time-series co-vary and discards any difference in magnitude, FC uses the correlation distance to quantify functional coupling.

#### Averaging model weights across temporal delays

To account for the delayed hemodynamic response function, each stimulus- or task-related feature is concatenated across four temporal delays (2 s, 4 s, 6 s, and 8 s as described above in *Feature adjustments*). Each feature is therefore associated with four model weights that reflect how well the feature value corresponds to the BOLD response at the time of the stimulus occurrence, 2 seconds after occurrence, 4 seconds after occurrence, and 8 seconds after occurrence. To recover a single model weight for each feature, the model weights were averaged across delays.

#### Re-scaling model weights

MC is based on model weights estimated with regularized linear regression. To optimize model prediction accuracy within each vertex, the regularization hyperparameter is allowed to vary across vertices, affecting the scale of the model weights. Because MC uses the Euclidean distance between model weights to recover networks, any difference in scale across vertices that is not corrected for could potentially bias estimates of the similarity of model weights. For example, vertices with a similar regularization hyperparameter will have similar scales of weights, and they may be grouped together even if they do not share similar functional tuning.

A common solution to address differences in the scale of the model weights is to use a global regularization hyperparameter that is optimized across all vertices^40^. Using a global regularization hyperparameter therefore ensures that all weights have a similar scale. However, because the hyperparameter is not optimized separately for each vertex, this approach leads to worse model prediction accuracy. Thus, to maximize prediction accuracy and to prevent biased estimates from differences in the scale of the model weights, we optimized the regularization hyperparameter for each vertex, and implemented an additional step that adjusts the scale of the model weights. The model weights in each vertex were first scaled to the unit norm, and then re-scaled by the cross-validated prediction accuracy in the training set (negative prediction accuracies were set to zero). This re-scaling step effectively corrects for differences in the scale of the model weights, while maximizing model prediction accuracy within each vertex (see **Supplemental Figure 8**).

### Selecting the optimal number of networks

Hierarchical clustering was used to assign each vertex of the template surface to one of *k* groups. Each group of vertices defines a brain network, and the set of all *k* brain networks defines a *clustering solution.* If *k* was arbitrarily selected, then the recovered networks may not reflect the true patterns of functional coupling observed across the brain. Thus, to select the optimal *k*, and consequently the optimal set of brain networks, each clustering solution was evaluated according to how well the networks in the solution predicted brain activity across participants.

In a leave-one-participant-out cross-validation scheme, the model weights from *N*-1 participants were used to predict the held-out participant’s BOLD responses in each network of each clustering solution. First, clustering solutions with all possible numbers of networks (from 2 to 20,484 networks) were generated using data averaged across all possible sets of *N*-1 participants (9 sets of *N*-1 subjects). The MC clustering solutions were generated from the model weights averaged across *N*-1 participants, and the FC clustering solutions were generated from the training set BOLD responses averaged across *N*-1 participants.

Next, prediction accuracy was evaluated for all possible MC and FC clustering solutions (20,482 numbers of networks x 9 sets of *N*-1 subjects x 2 connectivity measures = 368,676 total solutions). The *N*-1 group-average semantic model weights were first averaged across the vertices within each network. The network-average model weights were then used to generate a prediction of the BOLD responses in the test set (we refer to this model prediction as the *network-predicted activity*; the network-predicted activity is the same for all vertices within a network). The prediction accuracy of each network was computed as the average correlation between the network-predicted activity and the held-out participant’s observed BOLD responses across all vertices within the network. The prediction accuracy of each clustering solution was computed as the average prediction accuracy across all networks in the clustering solution. This process was repeated for all *N* held-out participants, and the prediction accuracies of each clustering solution were averaged across participants.

Note that this framework uses encoding model weights to identify the optimal number of networks. In the present study, semantic encoding models were constructed to perform MC. The resulting encoding model weights were then used to evaluate prediction accuracy across all possible MC and FC clustering solutions. Conventional FC studies, however, do not model the functional data. Therefore, studies that assess FC alone cannot use this approach to identify the optimal number of networks.

The leave-one-participant-out cross-validation procedure described above was used to identify the optimal number of MC and FC networks that can be recovered in this dataset. To recover the final sets of MC and FC networks, the semantic model weights and BOLD time-series were averaged across all nine participants. The group-average model weights were used to recover the final group MC solution with the optimal number of MC networks, and the group-average BOLD time-series were used to recover the final group FC solution with the optimal number of FC networks.

### Assessing the prediction accuracy of each network

Evaluating the prediction accuracy of all possible MC and FC clustering solutions identifies the sets of MC and FC networks that best capture functional organization during the task. These optimal solutions maximize the average cross-participant prediction accuracy across networks within the solution. However, because prediction accuracy was maximized across networks, individual networks within the optimal solutions may not accurately reflect the underlying functional activity. We therefore computed the prediction accuracy of each individual network in the final MC and FC solutions to determine which networks accurately capture the underlying functional responses during the task.

To compute the prediction accuracy of each network in each participant, both the *N*-1 group-average semantic model weights and the held-out participant’s BOLD responses in the test set were averaged across vertices within the network. We refer to the weights averaged across vertices within a network as *network-average model weights*; similarly, we refer to the BOLD responses averaged across vertices within a network as *network-average BOLD responses*. The *N*-1 group’s network-average model weights were then used to predict the held-out participant’s network-average BOLD responses. This process was repeated for each participant. Finally, the prediction accuracy of a network was computed as the average prediction accuracy across participants.

Significance was evaluated for each network in each participant by comparing the *N*-1 group’s network-predicted activity to a null distribution ^40, 42^. To generate the null distribution, the network-predicted activity was first split into blocks of 10 TRs and randomly shuffled across time. The prediction accuracy for each permutation was computed as the correlation between the shuffled network-predicted activity and the participant’s network-average BOLD response in the test set. This process was repeated 10,000 times for each participant. The empirical p-value was computed as the proportion of the 10,000 permutations whose prediction accuracy exceeded the observed prediction accuracy. All p-values reported in this work were corrected for multiple comparisons using the Benjamini-Hochberg false discovery rate (FDR) procedure^115^.

### Assessing the signal quality of the fMRI data in each network

The temporal signal-to-noise ratio (tSNR) was used to assess the quality of the functional data recorded from different brain areas. Temporal SNR is computed as the temporal mean divided by the temporal standard deviation of the recorded BOLD responses, and quantifies the fMRI signal stability over time^54, 55^. Data from the two presentations of the test stimulus were used to compute two tSNR maps for each participant. The two tSNR maps were then averaged to create a single tSNR map for each participant. The group tSNR was computed by averaging the tSNR maps across participants. Finally, to evaluate the signal quality of each network in the MC and FC solutions, the group-average tSNR was averaged across all vertices within each network.

#### Comparing vertex-wise and network-average variability in tSNR

Because MC groups vertices according to their functional tuning, we hypothesized that the networks recovered by MC will not be confounded by spatial variability in SNR. In contrast, because FC groups vertices according to their BOLD responses, we hypothesized that the networks recovered by FC will be confounded by spatial variability in SNR. To test these hypotheses, we computed the correlation across vertices between the vertex-wise tSNR and the network-average tSNR. The correlation between vertex-wise and network-average tSNR indicates whether the spatial variability in tSNR is captured by the MC or FC solutions. A high correlation indicates that the network-average tSNR captures spatial variability in vertex-wise tSNR, and a correlation that is close to zero indicates that the network-average tSNR does not account for spatial differences in tSNR.

Significance was evaluated for the MC and FC solutions by comparing the observed correlation between the vertex-wise and network-average tSNR to a null distribution. A separate null distribution was generated for the MC and FC solutions, by generating random rotations of each clustering solution^50, 116^. First, a random three-dimensional rotation was generated. Then, this rotation was separately applied to the three-dimensional coordinates of the inflated, spherical surfaces of the left and right hemisphere ({freesurfer_subject}/surf/{hemi}.sphere from FreeSurfer). This process generates a randomly rotated version of the original clustering solution that preserves the local spatial relationships between vertices, the approximate relative sizes of the networks, and the symmetry across hemispheres. Temporal SNR was then averaged across vertices within the randomly generated networks, and the correlation between the vertex-wise and random network-average tSNR was computed. This process was repeated 10,000 times for both the MC solution and the FC solution. The empirical, FDR-corrected p-value was computed as the proportion of the 10,000 permutations whose correlation between the vertex-wise and the randomly rotated network-average tSNR exceeded the observed correlation.

### Interpreting the function of the recovered networks

*Matching the cortical distribution of the recovered networks to a canonical set of resting state networks*.

The function of each network recovered in the present work was estimated by comparing the cortical distribution of the network to a set of 17 networks defined by Yeo et al.^32^ (here we refer to these networks as the *Yeo networks*). For each network in the optimal MC and FC solutions, the Dice coefficient was first computed between the network and each of the 17 Yeo networks. The Yeo network with the highest Dice coefficient was then selected as the best match for the MC or FC network. Because the Yeo networks have already been functionally defined^32, 56^, the function assigned to the best matching Yeo network was used to estimate the function of each network recovered in the present study. To account for instances in which multiple MC or FC networks were matched to the same Yeo network, the MC and FC networks were grouped according to the function of their best matching Yeo network (e.g., two FC networks were matched to two canonical resting state attention networks), and the Dice coefficient was computed between the union of the group of networks and the union of the best matching Yeo networks. The resulting Dice coefficient indicates the degree to which the MC and FC solutions recover each of the canonical resting state networks.

Significance was evaluated between each group of MC and FC networks by comparing the observed Dice coefficient between the union of the group of networks and the union of the best matching Yeo networks to a null distribution. A separate null distribution was generated for each group of MC and FC networks, by first generating random rotations of the union of each group of networks (as described above in *Comparing the vertex-wise and network-average variability in tSNR*), and then computing the Dice coefficient between the randomly rotated networks and the union of the best matching Yeo networks. This process was repeated 10,000 times for each group of networks within the MC and FC solutions. The empirical, FDR-corrected p-value was computed as the proportion of the 10,000 permutations whose Dice coefficient exceeded the observed Dice coefficient.

#### Interpreting the semantic tuning of the networks recovered by model connectivity

The network-average model weights were used to interpret each network recovered by MC. These network-average model weights quantify the *semantic tuning* of each network, that is the degree to which brain activity in each network is modulated by the 985 semantic features recovered from the narrative auditory stimulus. The semantic tuning of each network was interpreted by taking the dot product between the semantic features extracted from the stimulus (10,470 words in the stimulus x 985 features) and the network-average weights (985 features). This process associates each of the 10,470 spoken words with a scalar value that indicates the degree to which the word is predicted to elicit a change in the BOLD activity within the network. To produce a high-level functional description for each network, the 100 words that were predicted to elicit the largest change in BOLD activity in the network were manually inspected and labeled.

#### Visualizing the semantic space

To visualize the semantic concepts that are represented by each network, a dimensionality reduction algorithm (UMAP)^117^ was used to project the semantic embeddings of all 10,470 spoken words onto a 2-dimensional space. This projection reduced the matrix *M* (10,470 x 985) to a matrix *∼M* (10,470 x 2), in which relatively more semantically similar words are relatively closer to one another. We refer to this 2-dimensional matrix *∼M* as the *semantic space*. To visualize the breadth of semantic concepts represented by each network, the 100 words that were predicted to elicit the largest BOLD response within the network were mapped to this semantic space.

Because UMAP is a stochastic algorithm, different instantiations of UMAP result in different semantic spaces. This instability largely depends on the initialization of the embedding, as UMAP iteratively updates this initial embedding to match the local similarity structure of the original data. Initializing UMAP with a 2-dimensional spectral embedding of the matrix M instead of a random initialization reduces the variability of the initialization, and consequently increases the stability of the resulting semantic space. Furthermore, because we used the semantic space purely for visualization purposes and did not interpret the precise distances between points in the space, the resulting interpretations do not depend on the precise parameters used to recover the semantic space.

### Recovering the group-level functional networks in individual participants

Two methods were used to map the group-level networks to each participant’s anatomy in order to preserve either the cortical distribution or the functional tuning of the group networks recovered by MC (see **Supplemental Figure 6** for a comparison between these two methods for each participant).

#### Method 1: preserving the cortical distribution of the group networks in individual participants

The first method preserves the cortical distribution of the group networks by using FreeSurfer to project the group MC solution to each participant’s cortical surface (we refer to the projection of the group networks to the participant’s surface as the *spatially mapped MC solution*). This method ensures that the cortical distribution of the MC networks is consistent across participants, but does not ensure that the functional tuning of each network is consistent across participants. Therefore, we assessed the consistency of the functional tuning of each group-level network in each participant. First, a semantic voxelwise encoding model was first fit to functional data in the participant’s native anatomical space. Second, pycortex was used to project the voxelwise model weights onto the participant’s cortical surface. Third, each participant’s vertex-wise model weights were then averaged across vertices within each network, quantifying the functional tuning of each group-level network in each participant. Lastly, to quantify the cross-participant consistency of functional tuning within each MC network, the network-average model weights from each individual participant were correlated across participants.

Significance of the cross-participant consistency of functional tuning within each network was evaluated by comparing the within-network and the between-network similarity of the model weights across participants. For each network in the MC group solution, the within-network similarities were computed by correlating the network-average model weights across participants (9 participants = 36 pairwise comparisons within each network), and the between-network similarities were computed by correlating the network-average model weights across networks and participants (9 participants, 12 networks = 4851 pairwise comparisons between networks). The significance of the difference between the average within-network similarity and the average between-network similarity was then evaluated for each MC network by comparing the observed difference to a null distribution. A separate null distribution was generated for each network by first randomly assigning 36 of the 4887 comparisons to the within-network group, and then assigning the remaining 4851 comparisons to the between-network group. The difference between the means of the two groups was then computed. This process was repeated 10,000 times for each network in the MC solution. The empirical, FDR-corrected p-value was computed as the proportion of the 10,000 permutations whose difference between the within- and between-network similarity exceeded the observed difference. A bootstrapping procedure was used to compute the 95% confidence intervals for the difference between the within-network and between-network similarity of each MC network. The bootstrapped distributions were computed by randomly sampling the within-network comparisons and the between-network comparisons with replacement. The 2.5 and 97.5 percentiles of the bootstrapped distribution provided the 95% confidence interval for the observed difference.

#### Method 2: preserving the functional tuning of the group networks in individual participants

The second method preserves the functional tuning of the group networks by assigning each vertex in each participant to the group MC network with the most similar functional tuning. First, a semantic voxelwise encoding model was fit to functional data in each participant’s native anatomical space. Second, pycortex was used to project both the voxelwise model weights and the voxelwise model prediction accuracy onto the participant’s cortical surface. Third, each significantly predicted vertex (*R*^2^ > 0.04; *p* < 0.05, permutation test) was assigned to one of the group MC networks by computing the correlation between the model weights of the vertex and the network-average model weights of each group MC network. The vertex was assigned to the MC network with the largest correlation value. This assignment procedure was repeated for all significantly predicted vertices, generating the set of participant-level MC networks that have the most similar functional tuning to the group MC networks (we refer to this set of participant-level networks as the *functionally matched MC solution*). This method ensures that the functional tuning of the MC networks is consistent across participants, but does not ensure that the cortical distribution of each network is consistent across participants. Therefore, the cross-participant spatial variability was assessed for each functionally matched network. To evaluate the cross-participant variability in cortical distribution, the Dice coefficient was computed between each network in the spatially mapped and functionally matched MC solutions.

### Data analysis implementation

The data analysis pipeline for Model Connectivity was implemented in Python (modelconn), and is publicly available [link will be provided here upon publication]. The package modelconn builds on the scientific Python ecosystem, using numpy^118^, matplotlib^119^, scipy^108^, scikit-learn^120^, umap-learn^117^, nipype^121^, nibabel^122^, pycortex^107^, cottoncandy^123^, and himalaya^44^ with a pytorch backend^124^.

## References

1. Bressler, S.L. (1995). Large-scale cortical networks and cognition. Brain Res. Rev. 20, 288–304. 10.1016/0165-0173(94)00016-i.

2. Goldman-Rakic, P.S. (1988). Topography of cognition: parallel distributed networks in primate association cortex. Annu. Rev. Neurosci. 11, 137–156. 10.1146/annurev.ne.11.030188.001033.

3. Mesulam, M.M. (1990). Large-scale neurocognitive networks and distributed processing for attention, language, and memory. Ann. Neurol. 28, 597–613. 10.1002/ana.410280502.

4. Young, M.P., Scannell, J.W., Burns, G.A., and Blakemore, C. (1994). Analysis of connectivity: neural systems in the cerebral cortex. Rev. Neurosci. 5, 227–249. 10.1515/revneuro.1994.5.3.227.

5. Birn, R.M. (2012). The role of physiological noise in resting-state functional connectivity. NeuroImage 62, 864–870. 10.1016/j.neuroimage.2012.01.016.

6. Biswal, B., Yetkin, F.Z., Haughton, V.M., and Hyde, J.S. (1995). Functional connectivity in the motor cortex of resting human brain using echo-planar MRI. Magn. Reson. Med. 34, 537–541. 10.1002/mrm.1910340409.

7. Lowe, M.J., Dzemidzic, M., Lurito, J.T., Mathews, V.P., and Phillips, M.D. (2000). Correlations in low-frequency BOLD fluctuations reflect cortico-cortical connections. NeuroImage 12, 582–587. 10.1006/nimg.2000.0654.

8. Cordes, D., Haughton, V.M., Arfanakis, K., Carew, J.D., Turski, P.A., Moritz, C.H., Quigley, M.A., and Elizabeth Meyerand, M. (2001). Frequencies contributing to functional connectivity in the cerebral cortex in “resting-state” data. AJNR Am. J. Neuroradiol. 22, 1326–1333.

9. Hampson, M., Peterson, B.S., Skudlarski, P., Gatenby, J.C., and Gore, J.C. (2002). Detection of functional connectivity using temporal correlations in MR images. Hum. Brain Mapp. 15, 247–262. 10.1002/hbm.10022.

10. Murphy, K., Birn, R.M., and Bandettini, P.A. (2013). Resting-state fMRI confounds and cleanup. NeuroImage 80, 349–359. 10.1016/j.neuroimage.2013.04.001.

11. Liu, T.T. (2016). Noise contributions to the fMRI signal: an overview. NeuroImage 143, 141–151. 10.1016/j.neuroimage.2016.09.008.

12. Mitra, P.P., Ogawa, S., Hu, X., and Uğurbil, K. (1997). The nature of spatiotemporal changes in cerebral hemodynamics as manifested in functional magnetic resonance imaging. Magn. Reson. Med. 37, 511–518. 10.1002/mrm.1910370407.

13. Gati, J.S., Menon, R.S., Ugurbil, K., and Rutt, B.K. (1997). Experimental determination of the BOLD field strength dependence in vessels and tissue. Magn. Reson. Med. 38, 296–302. 10.1002/mrm.1910380220.

14. Dagli, M.S., Ingeholm, J.E., and Haxby, J.V. (1999). Localization of cardiac-induced signal change in fMRI. NeuroImage 9, 407–415. 10.1006/nimg.1998.0424.

15. Handwerker, D.A., Gonzalez-Castillo, J., D’Esposito, M., and Bandettini, P.A. (2012). The continuing challenge of understanding and modeling hemodynamic variation in fMRI. NeuroImage 62, 1017–1023. 10.1016/j.neuroimage.2012.02.015.

16. Greve, D.N., Brown, G.G., Mueller, B.A., Glover, G., and Liu, T.T. (2013). A survey of the sources of noise in fMRI. Psychometrika 78, 396–416. 10.1007/s11336-012-9294-0.

17. Tong, Y., and Frederick, B.D. (2014). Studying the spatial distribution of physiological effects on BOLD signals using ultrafast fMRI. Front. Hum. Neurosci. 8, 196. 10.3389/fnhum.2014.00196.

18. Finn, E.S. (2021). Is it time to put rest to rest? Trends Cogn. Sci. 25, 1021–1032. 10.1016/j.tics.2021.09.005.

19. Buckner, R.L., Krienen, F.M., and Yeo, B.T.T. (2013). Opportunities and limitations of intrinsic functional connectivity MRI. Nat. Neurosci. 16, 832–837. 10.1038/nn.3423.

20. Vanderwal, T., Eilbott, J., Finn, E.S., Craddock, R.C., Turnbull, A., and Castellanos, F.X. (2017). Individual differences in functional connectivity during naturalistic viewing conditions. NeuroImage 157, 521–530. 10.1016/j.neuroimage.2017.06.027.

21. Zamani Esfahlani, F., Jo, Y., Faskowitz, J., Byrge, L., Kennedy, D.P., Sporns, O., and Betzel, R.F. (2020). High-amplitude cofluctuations in cortical activity drive functional connectivity. Proc. Natl. Acad. Sci. U.S.A. 117, 28393– 28401. 10.1073/pnas.2005531117.

22. Finn, E.S., and Bandettini, P.A. (2021). Movie-watching outperforms rest for functional connectivity-based prediction of behavior. Neuroimage 235, 117963. 10.1016/j.neuroimage.2021.117963.

23. Tian, L., Ye, M., Chen, C., Cao, X., and Shen, T. (2021). Consistency of functional connectivity across different movies. NeuroImage 233, 117926. 10.1016/j.neuroimage.2021.117926.

24. Gal, S., Coldham, Y., Tik, N., Bernstein-Eliav, M., and Tavor, I. (2022). Act natural: Functional connectivity from naturalistic stimuli fMRI outperforms resting-state in predicting brain activity. NeuroImage 258, 119359. 10.1016/j.neuroimage.2022.119359.

25. Greicius, M.D., Krasnow, B., Reiss, A.L., and Menon, V. (2003). Functional connectivity in the resting brain: a network analysis of the default mode hypothesis. Proc. Natl. Acad. Sci. U.S.A. 100, 253–258. 10.1073/pnas.0135058100.

26. Beckmann, C.F., DeLuca, M., Devlin, J.T., and Smith, S.M. (2005). Investigations into resting-state connectivity using independent component analysis. Philos. Trans. R. Soc. Lond. B Biol. Sci. 360, 1001–1013. 10.1098/rstb.2005.1634.

27. Fox, M.D., Corbetta, M., Snyder, A.Z., Vincent, J.L., and Raichle, M.E. (2006). Spontaneous neuronal activity distinguishes human dorsal and ventral attention systems. Proc. Natl. Acad. Sci. U.S.A. 103, 10046–10051. 10.1073/pnas.0604187103.

28. Dosenbach, N.U.F., Fair, D.A., Miezin, F.M., Cohen, A.L., Wenger, K.K., Dosenbach, R.A.T., Fox, M.D., Snyder, A.Z., Vincent, J.L., Raichle, M.E., et al. (2007). Distinct brain networks for adaptive and stable task control in humans. Proc. Natl. Acad. Sci. U.S.A. 104, 11073–11078. 10.1073/pnas.0704320104.

29. Seeley, W.W., Menon, V., Schatzberg, A.F., Keller, J., Glover, G.H., Kenna, H., Reiss, A.L., and Greicius, M.D. (2007). Dissociable intrinsic connectivity networks for salience processing and executive control. J. Neurosci. 27, 2349–2356. 10.1523/JNEUROSCI.5587-06.2007.

30. Vincent, J.L., Kahn, I., Snyder, A.Z., Raichle, M.E., and Buckner, R.L. (2008). Evidence for a frontoparietal control system revealed by intrinsic functional connectivity. J. Neurophysiol. 100, 3328–3342. 10.1152/jn.90355.2008.

31. Smith, S.M., Fox, P.T., Miller, K.L., Glahn, D.C., Fox, P.M., Mackay, C.E., Filippini, N., Watkins, K.E., Toro, R., Laird, A.R., et al. (2009). Correspondence of the brain’s functional architecture during activation and rest. Proc. Natl. Acad. Sci. U.S.A. 106, 13040–13045. 10.1073/pnas.0905267106.

32. Yeo, B.T.T., Krienen, F.M., Sepulcre, J., Sabuncu, M.R., Lashkari, D., Hollinshead, M., Roffman, J.L., Smoller, J.W., Zöllei, L., Polimeni, J.R., et al. (2011). The organization of the human cerebral cortex estimated by intrinsic functional connectivity. J. Neurophysiol. 106, 1125–1165. 10.1152/jn.00338.2011.

33. Power, J.D., Cohen, A.L., Nelson, S.M., Wig, G.S., Barnes, K.A., Church, J.A., Vogel, A.C., Laumann, T.O., Miezin, F.M., Schlaggar, B.L., et al. (2011). Functional network organization of the human brain. Neuron 72, 665– 678. 10.1016/j.neuron.2011.09.006.

34. Uddin, L.Q., Yeo, B.T.T., and Spreng, R.N. (2019). Towards a universal taxonomy of macro-scale functional human brain networks. Brain Topogr. 32, 926–942. 10.1007/s10548-019-00744-6.

35. Bijsterbosch, J., Harrison, S.J., Jbabdi, S., Woolrich, M., Beckmann, C., Smith, S., and Duff, E.P. (2020). Challenges and future directions for representations of functional brain organization. Nat. Neurosci. 23, 1484–1495. 10.1038/s41593-020-00726-z.

36. Wu, M.C.-K., David, S.V., and Gallant, J.L. (2006). Complete functional characterization of sensory neurons by system identification. Annu. Rev. Neurosci. 29, 477–505. 10.1146/annurev.neuro.29.051605.113024.

37. Naselaris, T., Kay, K.N., Nishimoto, S., and Gallant, J.L. (2011). Encoding and decoding in fMRI. NeuroImage 56, 400–410. 10.1016/j.neuroimage.2010.07.073.

38. Kay, K.N., Naselaris, T., Prenger, R.J., and Gallant, J.L. (2008). Identifying natural images from human brain activity. Nature 452, 352–355. 10.1038/nature06713.

39. Nishimoto, S., Vu, A.T., Naselaris, T., Benjamini, Y., Yu, B., and Gallant, J.L. (2011). Reconstructing visual experiences from brain activity evoked by natural movies. Curr. Biol. 21, 1641–1646. 10.1016/j.cub.2011.08.031.

40. Huth, A.G., de Heer, W.A., Griffiths, T.L., Theunissen, F.E., and Gallant, J.L. (2016). Natural speech reveals the semantic maps that tile human cerebral cortex. Nature 532, 453–458. 10.1038/nature17637.

41. Lescroart, M.D., and Gallant, J.L. (2019). Human scene-selective areas represent 3D configurations of surfaces. Neuron 101, 178–192.e7. 10.1016/j.neuron.2018.11.004.

42. Deniz, F., Nunez-Elizalde, A.O., Huth, A.G., and Gallant, J.L. (2019). The representation of semantic information across human cerebral cortex during listening versus reading is invariant to stimulus modality. J. Neurosci. 39, 7722– 7736. 10.1523/JNEUROSCI.0675-19.2019.

43. Nunez-Elizalde, A.O., Huth, A.G., and Gallant, J.L. (2019). Voxelwise encoding models with non-spherical multivariate normal priors. NeuroImage 197, 482–492. 10.1016/j.neuroimage.2019.04.012.

44. Dupré la Tour, T., Eickenberg, M., Nunez-Elizalde, A.O., and Gallant, J.L. (2022). Feature-space selection with banded ridge regression. NeuroImage 264, 119728. 10.1016/j.neuroimage.2022.119728.

45. Horwitz, B. (2003). The elusive concept of brain connectivity. NeuroImage 19, 466–470. 10.1016/s1053-8119(03)00112-5.

46. Reid, A.T., Headley, D.B., Mill, R.D., Sanchez-Romero, R., Uddin, L.Q., Marinazzo, D., Lurie, D.J., Valdés-Sosa, P.A., Hanson, S.J., Biswal, B.B., et al. (2019). Advancing functional connectivity research from association to causation. Nat. Neurosci. 22, 1751–1760. 10.1038/s41593-019-0510-4.

47. Binder, J.R., Desai, R.H., Graves, W.W., and Conant, L.L. (2009). Where is the semantic system? A critical review and meta-analysis of 120 functional neuroimaging studies. Cereb. Cortex 19, 2767–2796. 10.1093/cercor/bhp055.

48. Thirion, B., Flandin, G., Pinel, P., Roche, A., Ciuciu, P., and Poline, J.-B. (2006). Dealing with the shortcomings of spatial normalization: multi-subject parcellation of fMRI datasets. Hum. Brain Mapp. 27, 678–693. 10.1002/hbm.20210.

49. Craddock, R.C., James, G.A., Holtzheimer, P.E., 3rd, Hu, X.P., and Mayberg, H.S. (2012). A whole brain fMRI atlas generated via spatially constrained spectral clustering. Hum. Brain Mapp. 33, 1914–1928. 10.1002/hbm.21333.

50. Gordon, E.M., Laumann, T.O., Adeyemo, B., Huckins, J.F., Kelley, W.M., and Petersen, S.E. (2016). Generation and evaluation of a cortical area parcellation from resting-state correlations. Cereb. Cortex 26, 288–303. 10.1093/cercor/bhu239.

51. Schaefer, A., Kong, R., Gordon, E.M., Laumann, T.O., Zuo, X.-N., Holmes, A.J., Eickhoff, S.B., and Yeo, B.T.T. (2018). Local-global parcellation of the human cerebral cortex from intrinsic functional connectivity MRI. Cereb. Cortex 28, 3095–3114. 10.1093/cercor/bhx179.

52. Bellec, P., Rosa-Neto, P., Lyttelton, O.C., Benali, H., and Evans, A.C. (2010). Multi-level bootstrap analysis of stable clusters in resting-state fMRI. NeuroImage 51, 1126–1139. 10.1016/j.neuroimage.2010.02.082.

53. Thirion, B., Varoquaux, G., Dohmatob, E., and Poline, J.-B. (2014). Which fMRI clustering gives good brain parcellations? Front. Neurosci. 8, 167. 10.3389/fnins.2014.00167.

54. Murphy, K., Bodurka, J., and Bandettini, P.A. (2007). How long to scan? The relationship between fMRI temporal signal to noise ratio and necessary scan duration. NeuroImage 34, 565–574. 10.1016/j.neuroimage.2006.09.032.

55. Parrish, T.B., Gitelman, D.R., LaBar, K.S., and Mesulam, M.M. (2000). Impact of signal-to-noise on functional MRI. Magn. Reson. Med. 44, 925–932. 10.1002/1522-2594(200012)44:6<925::aid-mrm14>3.0.co;2-m.

56. Kong, R., Li, J., Orban, C., Sabuncu, M.R., Liu, H., Schaefer, A., Sun, N., Zuo, X.-N., Holmes, A.J., Eickhoff, S.B., et al. (2019). Spatial topography of individual-specific cortical networks predicts human cognition, personality, and emotion. Cereb. Cortex 29, 2533–2551. 10.1093/cercor/bhy123.

57. Laumann, T.O., Gordon, E.M., Adeyemo, B., Snyder, A.Z., Joo, S.J., Chen, M.-Y., Gilmore, A.W., McDermott, K.B., Nelson, S.M., Dosenbach, N.U.F., et al. (2015). Functional system and areal organization of a highly sampled individual human brain. Neuron 87, 657–670. 10.1016/j.neuron.2015.06.037.

58. Dubois, J., and Adolphs, R. (2016). Building a science of individual differences from fMRI. Trends Cogn. Sci. 20, 425–443. 10.1016/j.tics.2016.03.014.

59. Welvaert, M., and Rosseel, Y. (2013). On the definition of signal-to-noise ratio and contrast-to-noise ratio for fMRI data. PLoS One 8, e77089. 10.1371/journal.pone.0077089.

60. Van Dijk, K.R.A., Sabuncu, M.R., and Buckner, R.L. (2012). The influence of head motion on intrinsic functional connectivity MRI. NeuroImage 59, 431–438. 10.1016/j.neuroimage.2011.07.044.

61. Power, J.D., Barnes, K.A., Snyder, A.Z., Schlaggar, B.L., and Petersen, S.E. (2012). Spurious but systematic correlations in functional connectivity MRI networks arise from subject motion. NeuroImage 59, 2142–2154. 10.1016/j.neuroimage.2011.10.018.

62. Birn, R.M., Diamond, J.B., Smith, M.A., and Bandettini, P.A. (2006). Separating respiratory-variation-related fluctuations from neuronal-activity-related fluctuations in fMRI. NeuroImage 31, 1536–1548. 10.1016/j.neuroimage.2006.02.048.

63. Gratton, C., Laumann, T.O., Nielsen, A.N., Greene, D.J., Gordon, E.M., Gilmore, A.W., Nelson, S.M., Coalson, R.S., Snyder, A.Z., Schlaggar, B.L., et al. (2018). Functional brain networks are dominated by stable group and individual factors, not cognitive or daily variation. Neuron 98, 439–452.e5. 10.1016/j.neuron.2018.03.035.

64. Greicius, M. (2008). Resting-state functional connectivity in neuropsychiatric disorders. Curr. Opin. Neurol. 21, 424–430. 10.1097/WCO.0b013e328306f2c5.

65. Supekar, K., Menon, V., Rubin, D., Musen, M., and Greicius, M.D. (2008). Network analysis of intrinsic functional brain connectivity in Alzheimer’s disease. PLoS Comput. Biol. 4, e1000100. 10.1371/journal.pcbi.1000100.

66. Zhang, J., Wang, J., Wu, Q., Kuang, W., Huang, X., He, Y., and Gong, Q. (2011). Disrupted brain connectivity networks in drug-naive, first-episode major depressive disorder. Biol. Psychiatry 70, 334–342. 10.1016/j.biopsych.2011.05.018.

67. Fair, D.A., Nigg, J.T., Iyer, S., Bathula, D., Mills, K.L., Dosenbach, N.U.F., Schlaggar, B.L., Mennes, M., Gutman, D., Bangaru, S., et al. (2013). Distinct neural signatures detected for ADHD subtypes after controlling for micro-movements in resting state functional connectivity MRI data. Front. Syst. Neurosci. 6, 80. 10.3389/fnsys.2012.00080.

68. Baker, J.T., Holmes, A.J., Masters, G.A., Yeo, B.T.T., Krienen, F., Buckner, R.L., and Öngür, D. (2014). Disruption of cortical association networks in schizophrenia and psychotic bipolar disorder. JAMA Psychiatry 71, 109–118. 10.1001/jamapsychiatry.2013.3469.

69. Sha, Z., Wager, T.D., Mechelli, A., and He, Y. (2019). Common Dysfunction of Large-Scale Neurocognitive Networks Across Psychiatric Disorders. Biol. Psychiatry 85, 379–388. 10.1016/j.biopsych.2018.11.011.

70. Finn, E.S., Shen, X., Scheinost, D., Rosenberg, M.D., Huang, J., Chun, M.M., Papademetris, X., and Constable, R.T. (2015). Functional connectome fingerprinting: identifying individuals using patterns of brain connectivity. Nat. Neurosci. 18, 1664–1671. 10.1038/nn.4135.

71. Gordon, E.M., Laumann, T.O., Gilmore, A.W., Newbold, D.J., Greene, D.J., Berg, J.J., Ortega, M., Hoyt-Drazen, C., Gratton, C., Sun, H., et al. (2017). Precision functional mapping of individual human brains. Neuron 95, 791–807.e7. 10.1016/j.neuron.2017.07.011.

72. Seitzman, B.A., Gratton, C., Laumann, T.O., Gordon, E.M., Adeyemo, B., Dworetsky, A., Kraus, B.T., Gilmore, A.W., Berg, J.J., Ortega, M., et al. (2019). Trait-like variants in human functional brain networks. Proc. Natl. Acad. Sci. U.S.A. 116, 22851–22861. 10.1073/pnas.1902932116.

73. Krienen, F.M., Yeo, B.T.T., and Buckner, R.L. (2014). Reconfigurable task-dependent functional coupling modes cluster around a core functional architecture. Philos. Trans. R. Soc. Lond. B Biol. Sci. 369, 20130526. 10.1098/rstb.2013.0526.

74. Mueller, S., Wang, D., Fox, M.D., Thomas Yeo, B.T., Sepulcre, J., Sabuncu, M.R., Shafee, R., Lu, J., and Liu, H. (2013). Individual variability in functional connectivity architecture of the human brain. Neuron 77, 586–595. 10.1016/j.neuron.2012.12.028.

75. Wang, D., Buckner, R.L., Fox, M.D., Holt, D.J., Holmes, A.J., Stoecklein, S., Langs, G., Pan, R., Qian, T., Li, K., et al. (2015). Parcellating cortical functional networks in individuals. Nat. Neurosci. 18, 1853–1860. 10.1038/nn.4164.

76. Moeller, S., Nallasamy, N., Tsao, D.Y., and Freiwald, W.A. (2009). Functional connectivity of the macaque brain across stimulus and arousal states. J. Neurosci. 29, 5897–5909. 10.1523/JNEUROSCI.0220-09.2009.

77. Rosenberg, M.D., Finn, E.S., Scheinost, D., Papademetris, X., Shen, X., Constable, R.T., and Chun, M.M. (2016). A neuromarker of sustained attention from whole-brain functional connectivity. Nat. Neurosci. 19, 165–171. 10.1038/nn.4179.

78. Tagliazucchi, E., and van Someren, E.J.W. (2017). The large-scale functional connectivity correlates of consciousness and arousal during the healthy and pathological human sleep cycle. NeuroImage 160, 55–72. 10.1016/j.neuroimage.2017.06.026.

79. Çukur, T., Nishimoto, S., Huth, A.G., and Gallant, J.L. (2013). Attention during natural vision warps semantic representation across the human brain. Nat. Neurosci. 16, 763–770. 10.1038/nn.3381.

80. Mason, M.F., Norton, M.I., Van Horn, J.D., Wegner, D.M., Grafton, S.T., and Macrae, C.N. (2007). Wandering minds: the default network and stimulus-independent thought. Science 315, 393–395. 10.1126/science.1131295.

81. Andrews-Hanna, J.R. (2012). The brain’s default network and its adaptive role in internal mentation. Neuroscientist 18, 251–270. 10.1177/1073858411403316.

82. Andreasen, N.C., O’Leary, D.S., Cizadlo, T., Arndt, S., Rezai, K., Watkins, G.L., Ponto, L.L., and Hichwa, R.D. (1995). Remembering the past: two facets of episodic memory explored with positron emission tomography. Am. J. Psychiatry 152, 1576–1585. 10.1176/ajp.152.11.1576.

83. Wagner, A.D., Shannon, B.J., Kahn, I., and Buckner, R.L. (2005). Parietal lobe contributions to episodic memory retrieval. Trends Cogn. Sci. 9, 445–453. 10.1016/j.tics.2005.07.001.

84. Mars, R., Neubert, F.-X., Noonan, M., Sallet, J., Toni, I., and Rushworth, M. (2012). On the relationship between the “default mode network” and the “social brain.” Front. Hum. Neurosci. 6, 189. 10.3389/fnhum.2012.00189.

85. Li, W., Mai, X., and Liu, C. (2014). The default mode network and social understanding of others: what do brain connectivity studies tell us. Front. Hum. Neurosci. 8, 74. 10.3389/fnhum.2014.00074.

86. Buckner, R.L., Andrews-Hanna, J.R., and Schacter, D.L. (2008). The brain’s default network: anatomy, function, and relevance to disease. Ann. N. Y. Acad. Sci. 1124, 1–38. 10.1196/annals.1440.011.

87. Smallwood, J., Bernhardt, B.C., Leech, R., Bzdok, D., Jefferies, E., and Margulies, D.S. (2021). The default mode network in cognition: a topographical perspective. Nat. Rev. Neurosci. 22, 503–513. 10.1038/s41583-021-00474-4.

88. Binder, J.R., Frost, J.A., Hammeke, T.A., Bellgowan, P.S., Rao, S.M., and Cox, R.W. (1999). Conceptual processing during the conscious resting state. A functional MRI study. J. Cogn. Neurosci. 11, 80–95. 10.1162/089892999563265.

89. Humphreys, G.F., Hoffman, P., Visser, M., Binney, R.J., and Lambon Ralph, M.A. (2015). Establishing task- and modality-dependent dissociations between the semantic and default mode networks. Proc. Natl. Acad. Sci. U.S.A. 112, 7857–7862. 10.1073/pnas.1422760112.

90. Ralph, M.A.L., Jefferies, E., Patterson, K., and Rogers, T.T. (2017). The neural and computational bases of semantic cognition. Nat. Rev. Neurosci. 18, 42–55. 10.1038/nrn.2016.150.

91. Cole, M.W., Bassett, D.S., Power, J.D., Braver, T.S., and Petersen, S.E. (2014). Intrinsic and task-evoked network architectures of the human brain. Neuron 83, 238–251. 10.1016/j.neuron.2014.05.014.

92. Greene, A.S., Gao, S., Noble, S., Scheinost, D., and Constable, R.T. (2020). How tasks change whole-brain functional organization to reveal brain-phenotype relationships. Cell Rep. 32, 108066. 10.1016/j.celrep.2020.108066.

93. Salehi, M., Greene, A.S., Karbasi, A., Shen, X., Scheinost, D., and Constable, R.T. (2020). There is no single functional atlas even for a single individual: Functional parcel definitions change with task. NeuroImage 208, 116366. 10.1016/j.neuroimage.2019.116366.

94. Kraus, B.T., Perez, D., Ladwig, Z., Seitzman, B.A., Dworetsky, A., Petersen, S.E., and Gratton, C. (2021). Network variants are similar between task and rest states. NeuroImage 229, 117743. 10.1016/j.neuroimage.2021.117743.

95. Popham, S.F., Huth, A.G., Bilenko, N.Y., Deniz, F., Gao, J.S., Nunez-Elizalde, A.O., and Gallant, J.L. (2021). Visual and linguistic semantic representations are aligned at the border of human visual cortex. Nat. Neurosci. 24, 1628– 1636. 10.1038/s41593-021-00921-6.

96. Vul, E., Lashkari, D., Hsieh, P.-J., Golland, P., and Kanwisher, N. (2012). Data-driven functional clustering reveals dominance of face, place, and body selectivity in the ventral visual pathway. J. Neurophysiol. 108, 2306–2322. 10.1152/jn.00354.2011.

97. Tarhan, L., and Konkle, T. (2020). Sociality and interaction envelope organize visual action representations. Nat. Commun. 11, 3002. 10.1038/s41467-020-16846-w.

98. LeBel, A., Jain, S., and Huth, A.G. (2021). Voxelwise encoding models show that cerebellar language representations are highly conceptual. J. Neurosci. 41, 10341–10355. 10.1523/JNEUROSCI.0118-21.2021.

99. Jenkinson, M., Bannister, P., Brady, M., and Smith, S. (2002). Improved optimization for the robust and accurate linear registration and motion correction of brain images. Neuroimage 17, 825–841. 10.1016/s1053-8119(02)91132-8.

100. Jenkinson, M., and Smith, S. (2001). A global optimisation method for robust affine registration of brain images. Med. Image Anal. 5, 143–156. 10.1016/s1361-8415(01)00036-6.

101. Jenkinson, M., Beckmann, C.F., Behrens, T.E.J., Woolrich, M.W., and Smith, S.M. (2012). FSL. Neuroimage 62, 782–790. 10.1016/j.neuroimage.2011.09.015.

102. Greve, D.N., and Fischl, B. (2009). Accurate and robust brain image alignment using boundary-based registration. Neuroimage 48, 63–72. 10.1016/j.neuroimage.2009.06.060.

103. Avants, B.B., Tustison, N., and Song, G. (2009). Advanced normalization tools (ANTS). Insight J. 2, 1–35.

104. Behzadi, Y., Restom, K., Liau, J., and Liu, T.T. (2007). A component based noise correction method (CompCor) for BOLD and perfusion based fMRI. NeuroImage 37, 90–101. 10.1016/j.neuroimage.2007.04.042.

105. Glover, G.H., Li, T.Q., and Ress, D. (2000). Image-based method for retrospective correction of physiological motion effects in fMRI: RETROICOR. Magn. Reson. Med. 44, 162–167. 10.1002/1522-2594(200007)44:1<162::aid-mrm23>3.0.co;2-e.

106. Esteban, O., Markiewicz, C.J., Blair, R.W., Moodie, C.A., Isik, A.I., Erramuzpe, A., Kent, J.D., Goncalves, M., DuPre, E., Snyder, M., et al. (2019). fMRIPrep: a robust preprocessing pipeline for functional MRI. Nat. Methods 16, 111–116. 10.1038/s41592-018-0235-4.

107. Gao, J.S., Huth, A.G., Lescroart, M.D., and Gallant, J.L. (2015). Pycortex: an interactive surface visualizer for fMRI. Front. Neuroinform. 9, 23. 10.3389/fninf.2015.00023.

108. Virtanen, P., Gommers, R., Oliphant, T.E., Haberland, M., Reddy, T., Cournapeau, D., Burovski, E., Peterson, P., Weckesser, W., Bright, J., et al. (2020). SciPy 1.0: fundamental algorithms for scientific computing in Python. Nat. Methods 17, 261–272. 10.1038/s41592-019-0686-2.

109. Ward, J.H. (1963). Hierarchical grouping to optimize an objective function. J. Am. Stat. Assoc. 58, 236–244. 10.1080/01621459.1963.10500845.

110. van den Heuvel, M.P., and Hulshoff Pol, H.E. (2010). Exploring the brain network: A review on resting-state fMRI functional connectivity. Eur. Neuropsychopharmacol. 20, 519–534. 10.1016/j.euroneuro.2010.03.008.

111. Eickhoff, S.B., Yeo, B.T.T., and Genon, S. (2018). Imaging-based parcellations of the human brain. Nat. Rev. Neurosci. 19, 672–686. 10.1038/s41583-018-0071-7.

112. Wig, G.S., Schlaggar, B.L., and Petersen, S.E. (2011). Concepts and principles in the analysis of brain networks. Ann. N. Y. Acad. Sci. 1224, 126–146. 10.1111/j.1749-6632.2010.05947.x.

113. Shen, X., Tokoglu, F., Papademetris, X., and Constable, R.T. (2013). Groupwise whole-brain parcellation from resting-state fMRI data for network node identification. NeuroImage 82, 403–415. 10.1016/j.neuroimage.2013.05.081.

114. Bijsterbosch, J.D., Woolrich, M.W., Glasser, M.F., Robinson, E.C., Beckmann, C.F., Van Essen, D.C., Harrison, S.J., and Smith, S.M. (2018). The relationship between spatial configuration and functional connectivity of brain regions. Elife 7, e32992. 10.7554/eLife.32992.

115. Benjamini, Y., and Hochberg, Y. (1995). Controlling the false discovery rate: a practical and powerful approach to multiple testing. J. R. Stat. Soc. Series B Stat. Methodol. 57, 289–300. 10.1111/j.2517-6161.1995.tb02031.x.

116. Alexander-Bloch, A.F., Shou, H., Liu, S., Satterthwaite, T.D., Glahn, D.C., Shinohara, R.T., Vandekar, S.N., and Raznahan, A. (2018). On testing for spatial correspondence between maps of human brain structure and function. NeuroImage 178, 540–551. 10.1016/j.neuroimage.2018.05.070.

117. McInnes, L., Healy, J., and Melville, J. (2018). UMAP: Uniform Manifold Approximation and Projection for dimension reduction. arXiv [stat.ML].

118. Harris, C.R., Millman, K.J., van der Walt, S.J., Gommers, R., Virtanen, P., Cournapeau, D., Wieser, E., Taylor, J., Berg, S., Smith, N.J., et al. (2020). Array programming with NumPy. Nature 585, 357–362. 10.1038/s41586-020-2649-2.

119. Hunter, J.D. (2007). Matplotlib: a 2D graphics environment. 9, 90–95. 10.1109/MCSE.2007.55.

120. Pedregosa, F., Varoquaux, G., Gramfort, A., Michel, V., Thirion, B., Grisel, O., Blondel, M., Müller, A., Nothman, J., Louppe, G., et al. (2012). Scikit-learn: machine learning in python. J. Mach. Learn. Res. 12, 2825–2830.

121. Gorgolewski, K., Burns, C.D., Madison, C., Clark, D., Halchenko, Y.O., Waskom, M.L., and Ghosh, S.S. (2011). Nipype: a flexible, lightweight and extensible neuroimaging data processing framework in python. Front. Neuroinform. 5, 13. 10.3389/fninf.2011.00013.

122. Brett, M., Markiewicz, C.J., Hanke, M., Côté, M.A., Cipollini, B., McCarthy, P., Dorota, J., Chen, C.P., Halchenko, Y.O., and Cottar, M. (2020). nipy/nibabel: 3.2.1. Zenodo.

123. O. Nunez-Elizalde, A., S. Gao, J., Zhang, T., and L. Gallant, J. (2018). Cottoncandy: scientific python package for easy cloud storage. J. Open Source Softw. 3, 890. 10.21105/joss.00890.

124. Paszke, A., Gross, S., Massa, F., Lerer, A., Bradbury, J., Chanan, G., Killeen, T., Lin, Z., Gimelshein, N., Antiga, L., et al. (2019). Pytorch: an imperative style, high-performance deep learning library. In Proceedings of the 33rd International Conference on Neural Information Processing Systems 10.5555/3454287.3455008.

