## Supplemental Figures for "Model connectivity: leveraging the power of encoding models to overcome the limitations of functional connectivity"

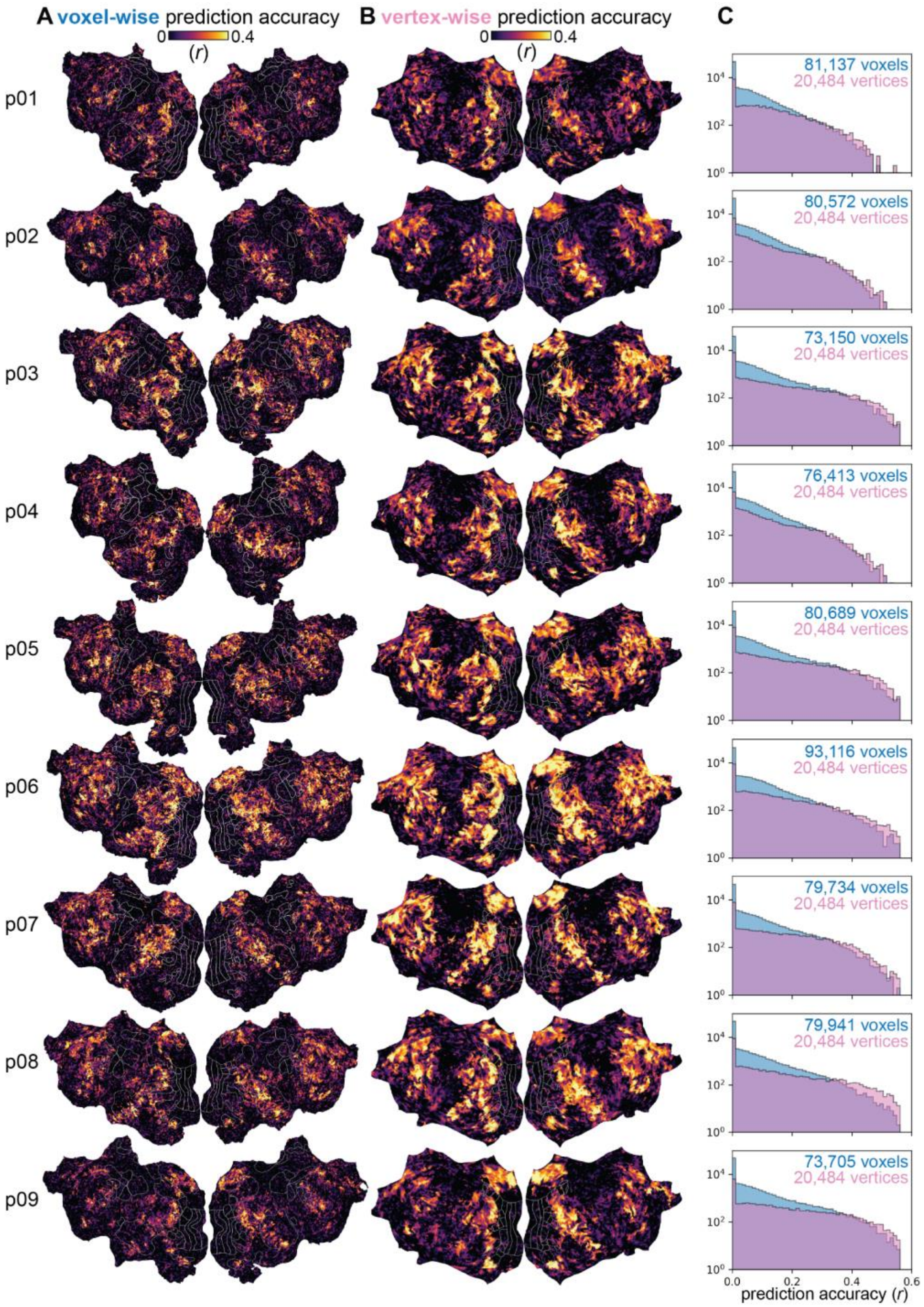


**Supplemental Figure 1. The spatial distribution of prediction accuracy within each participant is similar between the voxelwise encoding models fit in the participant's native space and the vertex-wise encoding models fit in the fsaverage5 template surface.** To recover a single set of networks that reflect patterns of functional coupling that generalize across participants, each individual participant's functional data were mapped to the fsaverage5 template surface. To assess how well the mapping preserves the functional organization that is unique to each participant, the spatial distribution of prediction accuracy within each participant was compared between two separate ridge regression models. In the first model, voxel-wise encoding models were fit to functional data measured from each cortical voxel in each participant's native anatomical space. In the second model, vertex-wise encoding models were fit to functional data that were mapped to the fsaverage5 template surface, which resulted in approximately a 3 mm FWHM smoothing. (A) For each participant, the voxel-wise prediction accuracy of the semantic encoding models is shown on the flattened surface of each participant (labeled as p01-09). (B) For each participant, the vertex-wise prediction accuracy of the semantic encoding models is shown on the flattened template surface. (C) The distribution of prediction accuracy across cortical voxels (colored in blue) and the distribution of prediction accuracy across template vertices (colored in pink) are plotted in a histogram. Prediction accuracies below zero are set to zero. The number of voxels in each participant is written in blue text, and the number of vertices in the template surface is written in pink text. Because the voxelwise models have more targets (voxels) than the vertex-wise models (difference = 59,344 ± 1,948 targets; mean ± s.e.m.), and because the vertex-wise models are fit to smoothed functional data, the voxelwise models tend to predict slightly worse than the vertex-wise models. However, the cortical regions that are well-predicted by the voxel-wise encoding models also appear to be well-predicted by the vertex-wise encoding models. This suggests that the mapping preserves the spatial distribution of prediction accuracy across the cortical surface for individual participants.

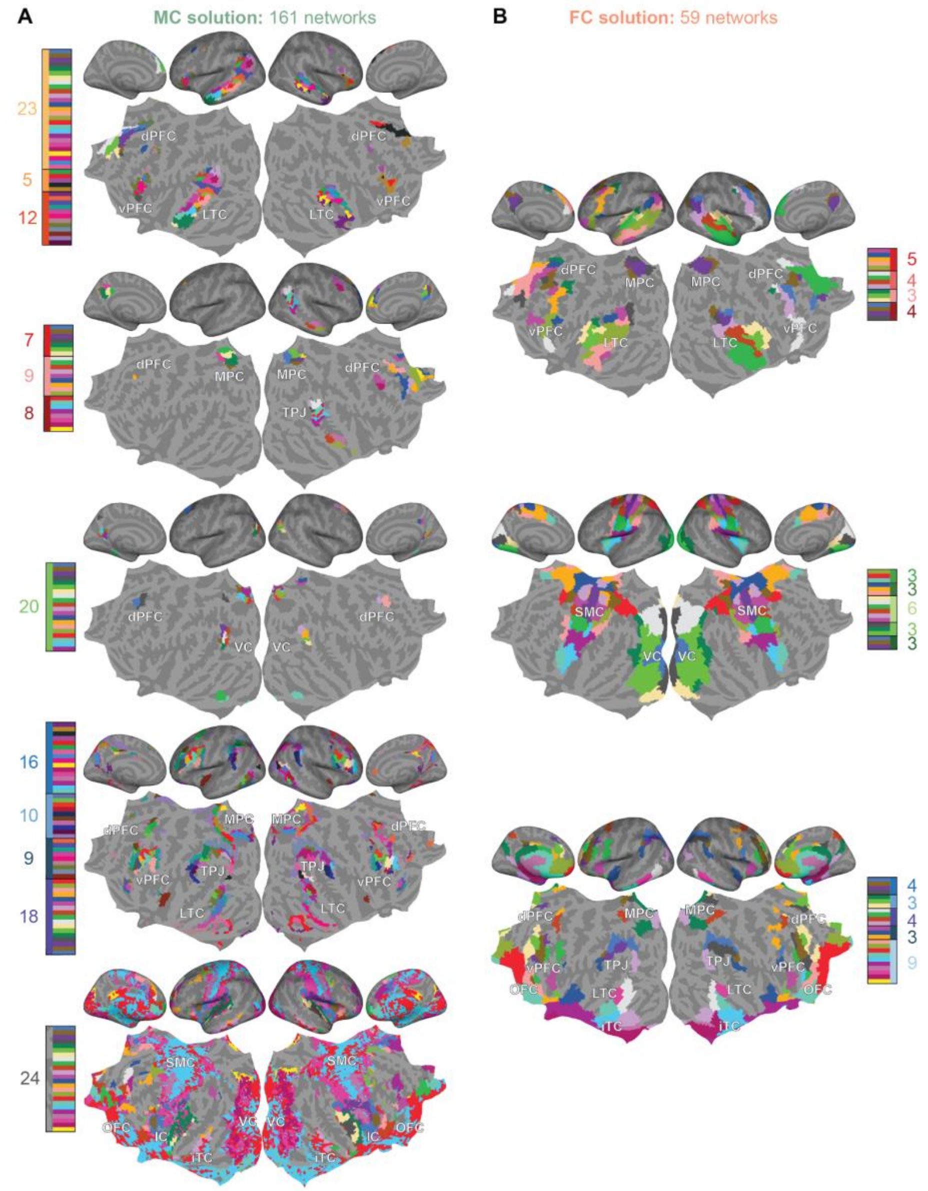


**Supplemental Figure 2. Visualizing the 161-network MC solution and the 59-network FC solution.** For the sake of exposition, the main text of the paper focuses on the 12-network MC solution and the 14-network FC solution, which corresponded to low-dimensional local maxima of the cross-participant prediction accuracy. However, the global maxima of the cross-participant prediction accuracy corresponds to the 161-network MC solution and the 59-network FC solution (see **Figure 2** in the main text). These solutions reflect the most spatially localized functional subdivisions that MC and FC can recover from the narrative auditory dataset. Here, we examine the functional organization recovered by these larger MC and FC solutions. Because hierarchical clustering was used to recover the clustering solutions, the 161-network MC solution corresponds to a finer subdivision of the 12-network MC solution, and the 59-network FC solution corresponds to a finer subdivision of the 14-network FC solution. To more easily visualize these large clustering solutions, (**A**) the 161-network MC solution was split into five groups of networks as in **Figure 3A** (colored in shades of orange, red, green, blue, and gray), and (**B**) the 59-network FC solution was split into three groups of networks as in **Figure 3D** (colored in shades of red, green, and blue). Each group of networks is shown on both the inflated and flattened template surface. The color bars indicate the colors that were randomly assigned to each of the 161 MC networks and to each of the 59 FC networks. The numbers indicate how many functional subdivisions exist in the higher-dimensional solution. For example, the MC network colored in green in the lower-dimensional solution is subdivided into 20 networks in the higher-dimensional solution. These analyses reveal that the 161-network MC solution primarily recovers functional subdivisions of areas involved in processing language. The first group of MC networks recovers 40 functional subdivisions across bilateral regions of the lateral temporal cortices (LTC), ventral prefrontal cortices (vPFC), and dorsal prefrontal cortices (dPFC). The second group of MC networks recovers 24 functional subdivisions across bilateral regions of the medial parietal cortices (MPC), and the LTC and dPFC in the right hemisphere. The third group of MC networks recovers 20 functional subdivisions across bilateral regions of higher order visual cortices (VC) and dPFC. The fourth group of MC networks recovers 62 functional subdivisions across bilateral regions of the vPFC, dPFC, and regions surrounding the MPC, temporal parietal junction (TPJ), and LTC. Finally, the fifth group of MC networks recovers 24 functional subdivisions across bilateral regions that are not well predicted by the semantic encoding models, such as the inferior temporal cortices (iTC), insular cortices (IC), orbitofrontal cortices (OFC), somatomotor cortices (SMC), and VC. By contrast, the 59-network FC solution recovers functional subdivisions across all regions of the cortical surface, regardless of their involvement in processing language. The first group of FC networks recovers 16 functional subdivisions across bilateral areas involved in processing language, including early auditory cortices, LTC, MPC, vPFC, and dPFC. The second group of FC networks recovers 18 functional subdivisions across bilateral areas not involved in processing language, including SMC and VC. Lastly, the third group of FC networks recovers 23 functional subdivisions across bilateral regions surrounding the MPC, TPJ, LTC, vPFC, and dPFC, and bilateral regions that suffer from susceptibility artifacts, including the OFC and iTC.

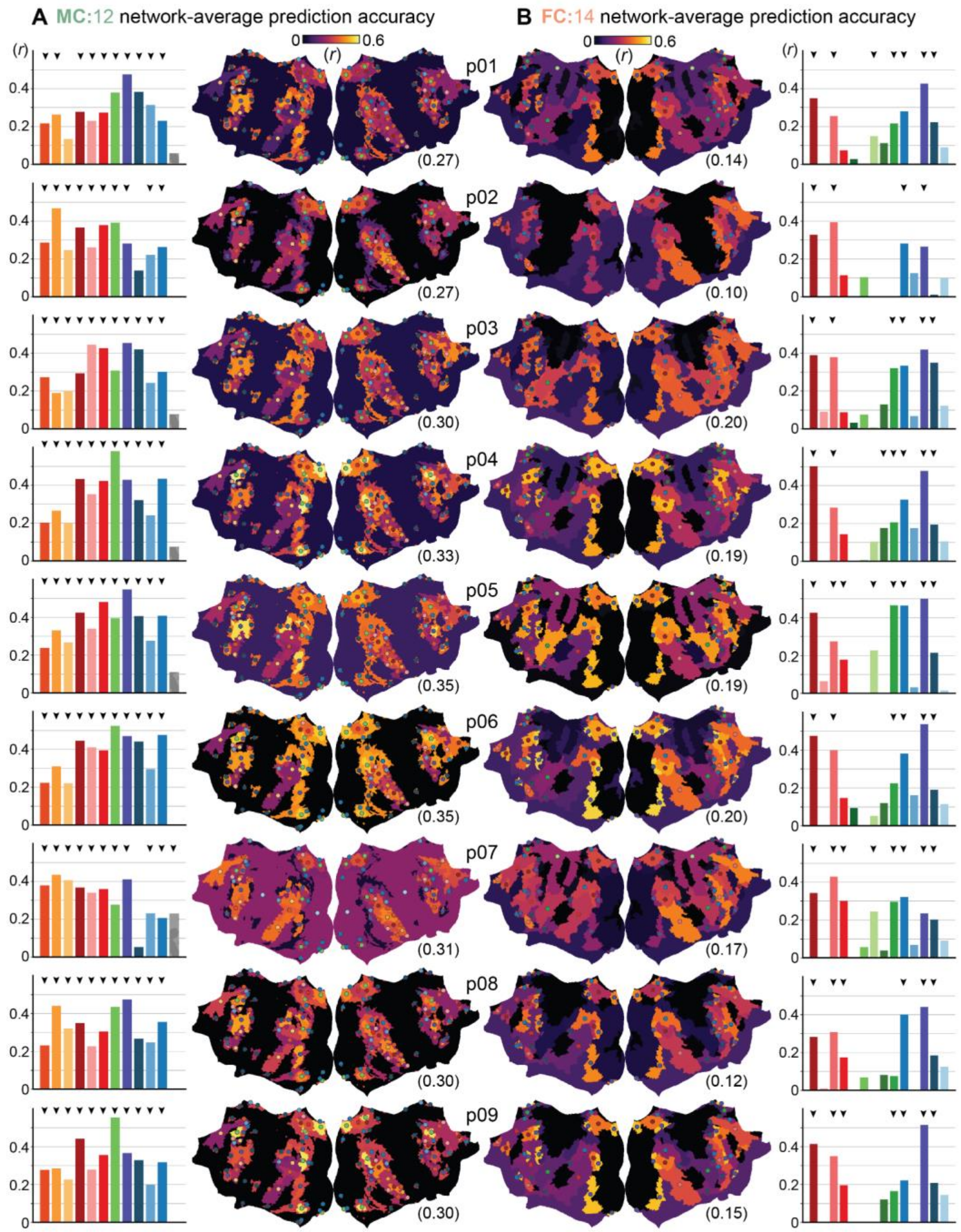


**Supplemental Figure 3. Prediction accuracy in each network in each participant.** The group-average semantic encoding models were used to predict network-average brain activity within each network of each participant. The resulting network-average prediction accuracies indicate how well the functional organization depicted by the 12-network MC solution and the 14-network FC solution generalizes to individual participants. (**A**, **B**) The prediction accuracy of each network in (**A**) the 12-network MC solution and (**B**) the 14-network FC solution is plotted separately for each participant in a bar plot and on the flattened template surface. In the bar plots, the color of each bar corresponds to the network assignment, and the arrows indicate the networks that significantly predict brain activity in each participant (at $p < 0.05$, permutation test). To convey the density of the significant functional subdivisions across the cortical surface, the colored circles on the flatmaps indicate the spatial centroids of the significantly predicted networks in each participant. The average prediction accuracy across all networks in each participant is indicated in parentheses. Of the 12 MC networks, nine networks significantly predict brain activity in all nine participants (networks colored in dark orange, orange, dark red, pink, red, medium green, dark purple, medium blue, and blue). The network colored in light orange significantly predicts brain activity in eight of the nine participants (in all but p01), the network colored in dark blue significantly predicts brain activity in seven of the nine participants (in all but p02 and p07), and the noise network significantly predicts brain activity in only one of the nine participants (in p07). Of the 14 FC networks, four networks significantly predict brain activity in all nine of the participants (networks colored in dark red, dark pink, blue, and dark purple). The network colored in dark blue significantly predicts activity in eight of the nine participants (in all but p02), the network colored in green significantly predicts activity in seven of the nine participants (in all but p02 and p08), the network colored in red significantly predicts activity in four of the nine participants (in p05, p07, p08, and p09), the network colored in light green significantly predicts activity in three of the nine participants (in p01, p05, and p07), and the network colored in dark green significantly predicts activity in one participant (in p04). The remaining networks colored in light pink, dark green, medium green, medium blue, and light blue do not significantly predict activity in any participant. These results indicate that the 12-network MC solution is a more accurate depiction of the functional organization of the semantic processing systems in individual participants than is the 14-network FC solution.

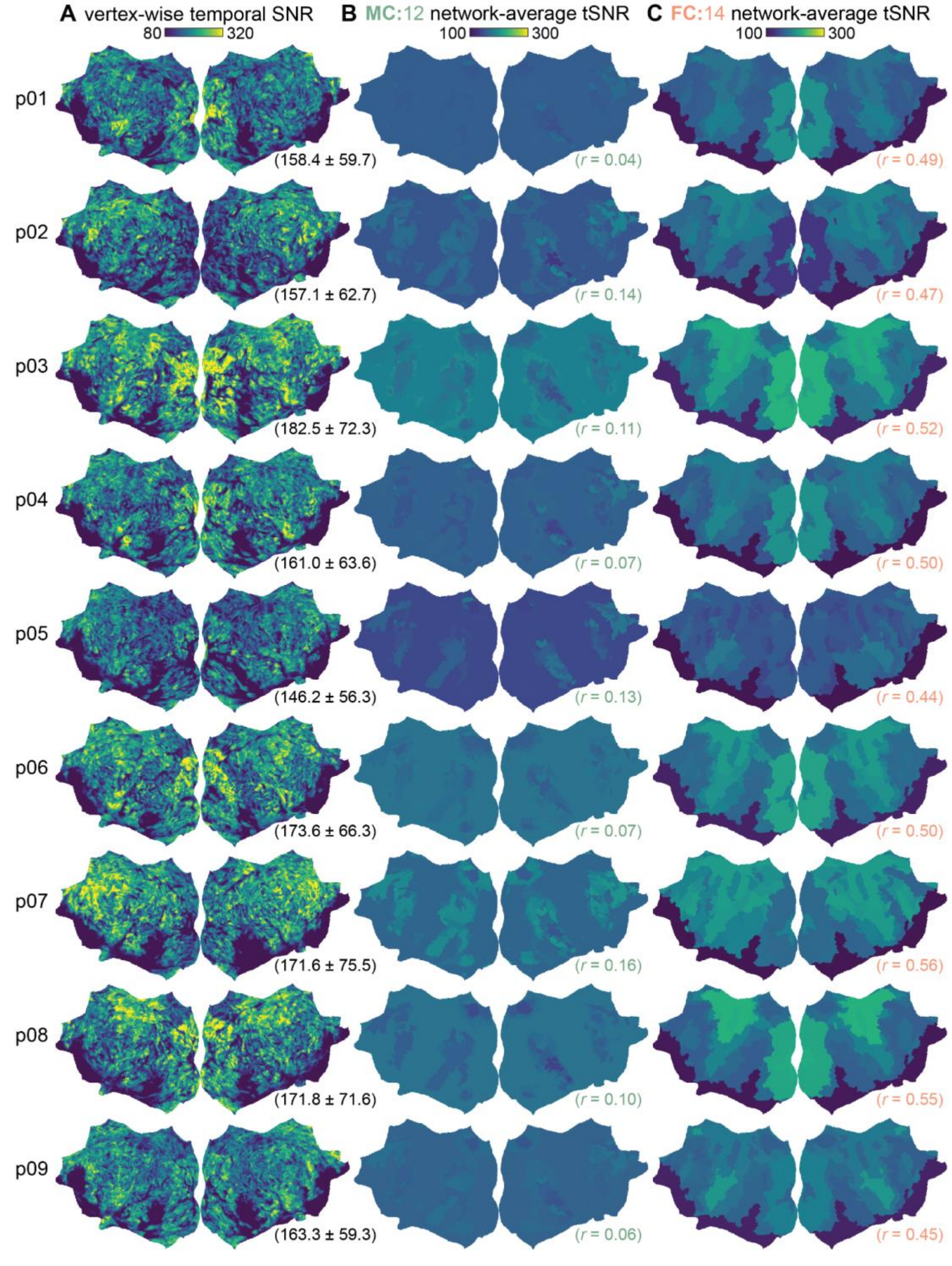


**Supplemental Figure 4. The networks recovered by FC are spatially correlated with the vertex-wise temporal SNR in each individual participant.** At the group level, the 12-network MC solution is not spatially correlated with the vertex-wise variability in tSNR, whereas the 14-network FC solution is spatially correlated with vertex-wise variability in tSNR. This indicates that only FC is confounded by spatial variability in tSNR. To assess whether this trend holds for individual participants, each participant's functional data was first used to compute the participant's tSNR. The network-average tSNR was then computed for both the 12-network MC and the 14-network FC solutions. Finally, for each participant, the participant's vertex-wise tSNR was correlated with the participant's network-average tSNR for both the MC and FC solutions. (**A**) Each participant's vertex-wise tSNR is shown on the flattened template surface. The mean and standard deviation of the vertex-wise tSNR is indicated in parentheses for each participant. While each participant has a distinct pattern of tSNR across their cortical surface, tSNR is consistently low in inferior temporal regions located near the ear canals, in inferior occipital regions, and at the center of the occipital pole. (**B**, **C**) Each participant's network-average tSNR is shown on the flattened template surface for each network in the (**B**) 12-network MC solution and (**C**) 14-network FC solution. The correlation between the participant's vertex-wise and network-average tSNR is indicated in parentheses. Across participants, the MC solution is not highly correlated with the participant's vertex-wise tSNR ($r = 0.10 \pm 0.01$; mean ± s.e.m.). By contrast, the FC solution is correlated with each participant's vertex-wise tSNR ($r = 0.50 \pm 0.01$). Thus, the networks recovered by MC do not reflect spatial variability in each participant's tSNR, while the networks recovered by FC capture spatial variability in each participant's tSNR.

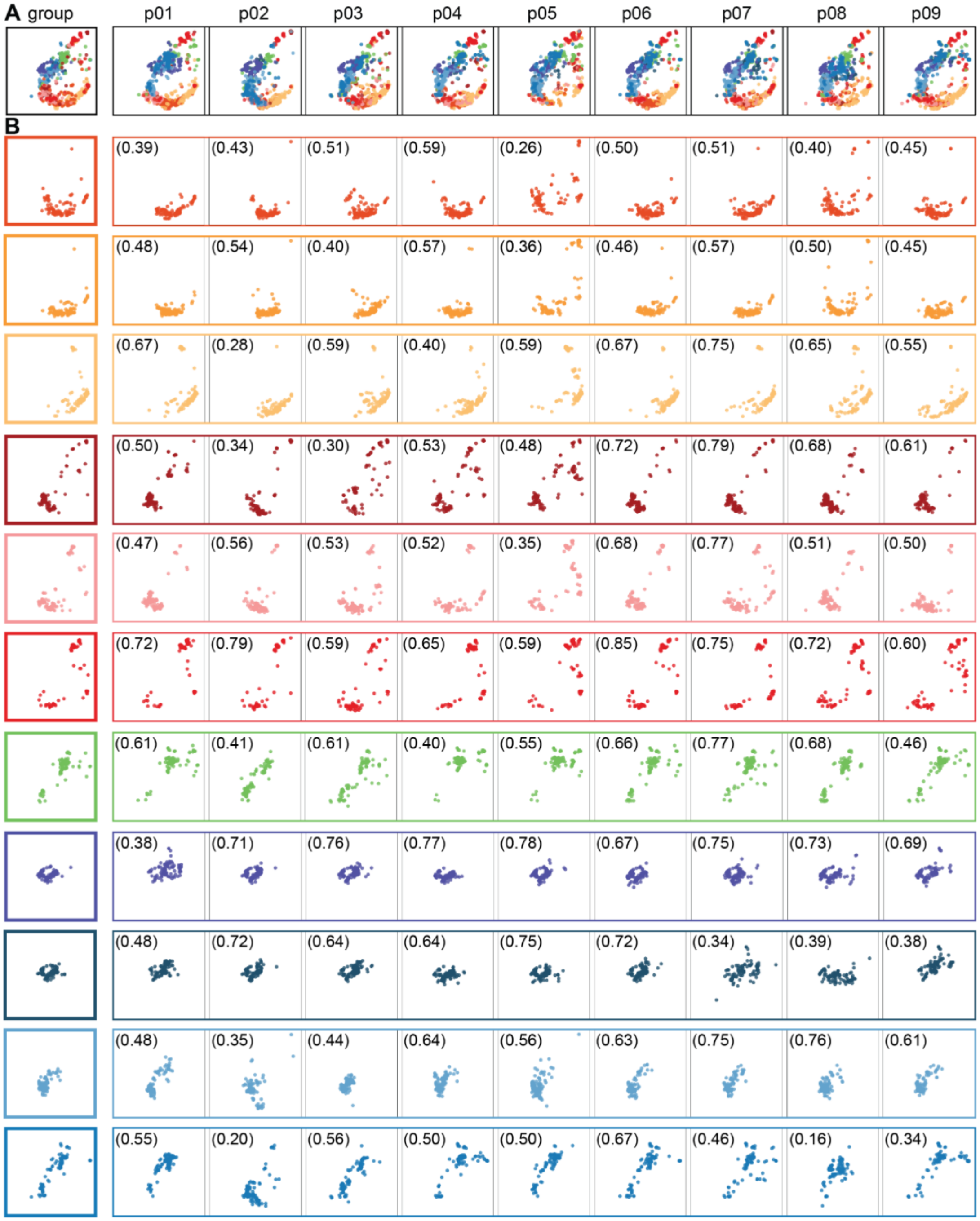


**Supplemental Figure 5. The semantic tuning within each MC network is largely consistent across individual participants.** The semantic model weights quantify the semantic information that is represented in the brain activity of a given network. However, because the group-average model weights were used to recover the 12 MC networks, the functional tuning of the network at the group level might not reflect the functional tuning of the network in individual participants. To assess the similarity between the group-level functional tuning of each network and the participant-level functional tuning of each network, we compared the 100 words that the group-level models predicted to elicit the largest BOLD response within each network, and the 100 words that the participant-level models predicted to elicit the largest BOLD response within each network. (**A**) The semantic coverage of the 11 semantic networks is plotted in a two-dimensional semantic space (as in **Figure 6B**). The group-average model weights were used to recover the group-level semantic coverage (labeled as group), and each participant's model weights were used to recover the participant-level semantic coverage (labeled as p01-09). Across participants, the set of 11 semantic networks has a similar semantic coverage. This suggests that the range of semantic concepts represented by the group-level networks accurately reflects the range of semantic concepts represented by the networks in individual participants. (**B**) To better visualize the semantic tuning of individual networks, the semantic coverage is plotted separately for each of the 11 semantic networks. The numbers indicate the proportion of the 100 words that both the group-level model and the participant-level model predict will elicit the largest BOLD response within each network. Plotting each network separately reveals that the semantic tuning within individual networks is largely consistent across participants.

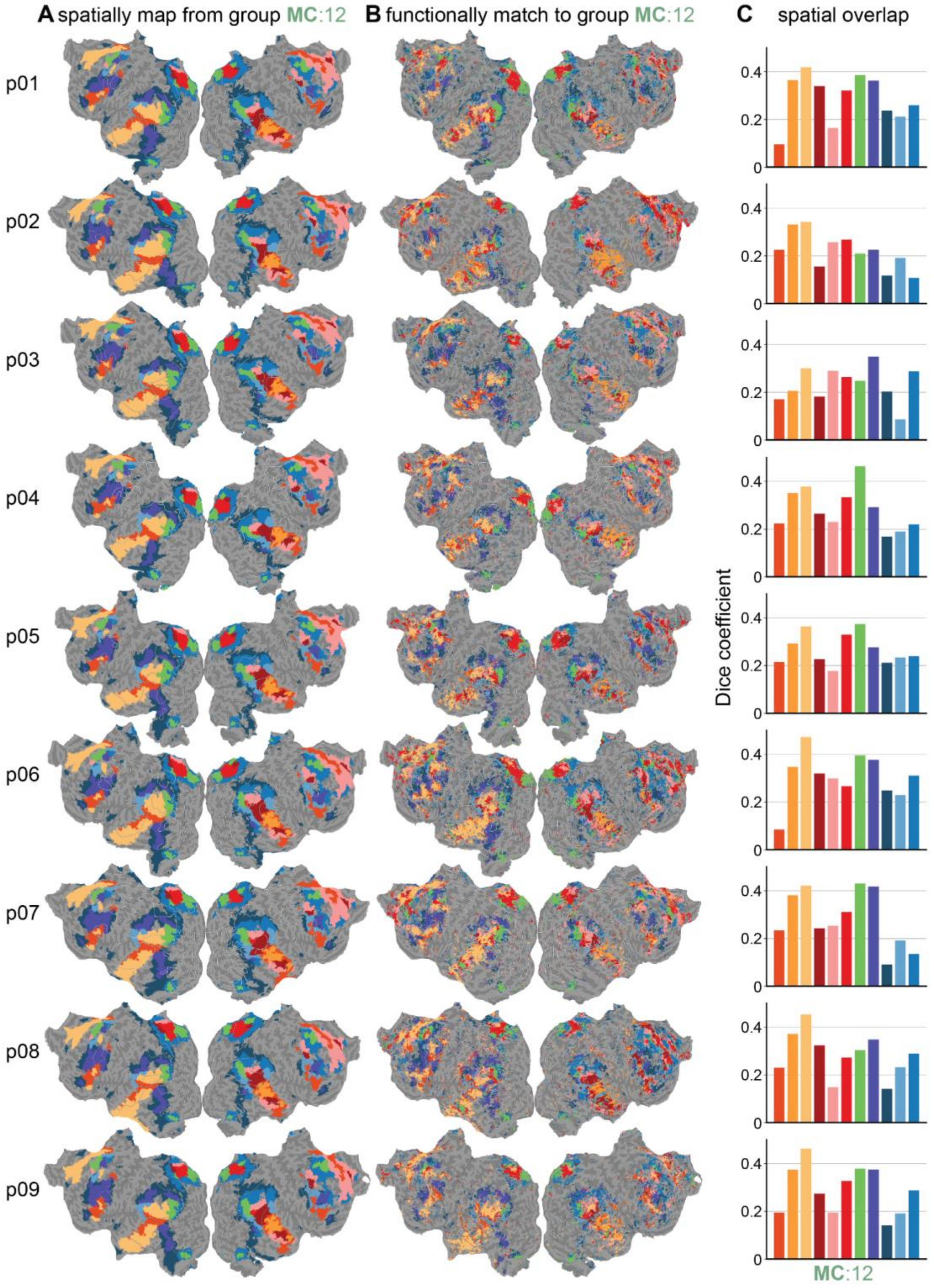


**Supplemental Figure 6. Comparison of the two methods that were used to recover the group MC networks in individual participants.** (**A**) The first method preserves the cortical distribution of each network in the group MC solution by using FreeSurfer to project the group MC solution from the fsaverage template surface to each participant's cortical surface. The resulting spatially mapped MC solution is shown for each participant on their flattened cortical surface. (**B**) The second method preserves the functional tuning of each network in the group MC solution by assigning each vertex of each participant's cortical surface to the group MC network with the most similar functional tuning. The resulting functionally matched MC solution is shown for each participant on their flattened cortical surface. (**C**) The spatial overlap between the spatially mapped and functionally matched MC networks is plotted in a bar graph for each participant. Spatial overlap was computed as the Dice coefficient between the group- and participant-level networks. Visually inspecting the spatially mapped and functionally matched MC solution in each participant reveals that the spatially mapped MC solution appears to capture some individual variability in the spatial organization of the group MC networks. For instance, in the vPFC of the left hemisphere, the network colored in dark purple has a slightly different shape and location, reflecting individual differences across participants. However, the functionally matched solutions also reveal regions in which the spatially mapped solution does not reflect the individual-specific patterns of functional organization. For instance, in the dPFC of the right hemisphere, the single network colored in light pink spans a large portion of the region at the group-level. By contrast, at the participant-level, this network is subdivided into multiple networks that do not appear to have consistent spatial organization across participants. Thus, this approach recovers individual differences in functional organization that are not consistent across participants.

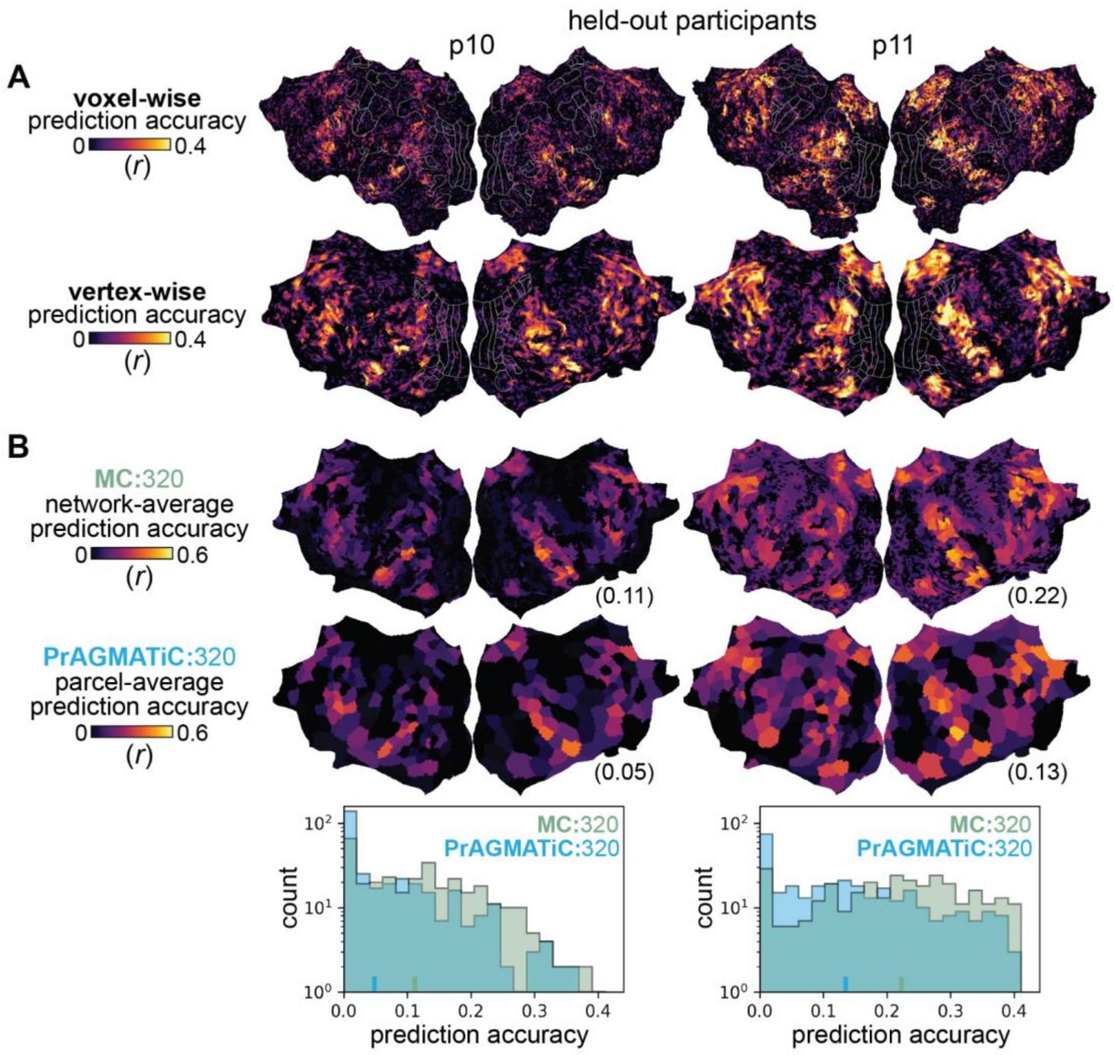


**Supplemental Figure 7. MC recovers networks that predict brain activity in held-out individuals better than PrAGMATiC.** The original publication of this dataset used a complex, Bayesian approach called PrAGMATiC to recover a set of parcels that depict the functionally homogeneous brain regions underlying natural language comprehension[40]. By contrast, the current work uses a simple, clustering approach called model connectivity to recover a set of networks that depict the functional organization of the brain during natural language comprehension. To compare these two approaches, we evaluated the prediction accuracy of the group MC solution and the group PrAGMATiC solution in two held-out participants. Data from these participants were not used to recover either the group MC nor the group PrAGMATiC solutions. (**A**) For both held-out participants (labeled as p10 and p11), the prediction accuracy of the participant's voxelwise semantic encoding models is shown on each participant's flattened cortical surface, and the prediction accuracy of the participant's vertex-wise semantic encoding models is shown on the flattened template surface. The brain activity in participant 11 is better predicted by the semantic encoding models than is the brain activity in participant 10. (**B**) To determine whether MC or PrAGMATiC recovers networks that better generalize across individuals, we evaluated the prediction accuracy of each network in the group MC solution and each parcel in the group PrAGMATiC solution in the two held-out participants. For ease of comparison, the number of networks was matched between the group MC and PrAGMATiC solutions. Prediction accuracy of the two group solutions in the two held-out participants is shown on the flattened template surface. The average prediction accuracy across networks and across parcels is indicated in parentheses. Below, the distribution of prediction accuracy across MC networks (colored in green) and the distribution of prediction accuracy across PrAGMATiC parcels (colored in blue) are plotted in a histogram. Prediction accuracies below zero are set to zero. Even though 320 is not the optimal number of networks that can be recovered by MC from this dataset, these 320 MC networks still predict brain activity in held-out participants better than the 320 PrAGMATiC parcels. This is likely due to the fact that PrAGMATiC parcellates the entirety of the cortical surface, whereas MC focuses on areas that are well predicted by the encoding models.

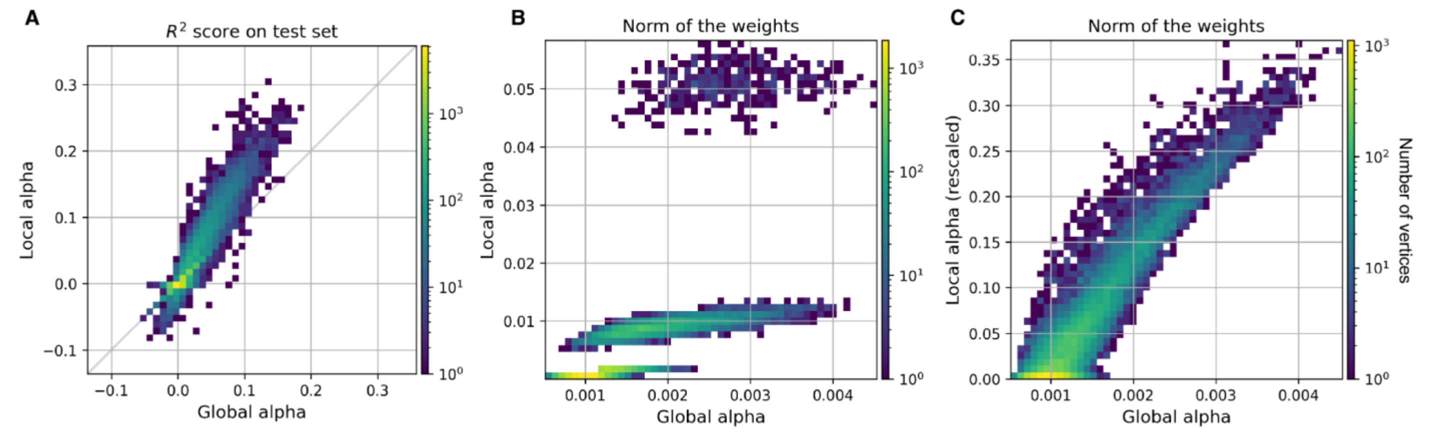


**Supplemental Figure 8. Re-scaling the model weights according to prediction accuracy in the training set corrects for vertex-wise differences in scale.** Two separate ridge regression models were estimated on the same data. In the first model, a single regularization hyperparameter was shared across all vertices (we refer to this model as the *global alpha* model); in the second model, the regularization hyperparameter was chosen independently for each vertex (we refer to this model as the *local alpha* model). All panels show a two-dimensional histogram with the global alpha model on the *x*-axis, and the local alpha model on the *y*-axis. (**A**) The model prediction accuracy on the test set is compared between the two models. Because the local alpha model optimizes the regularization hyperparameter independently for each vertex, the local alpha model provides better prediction accuracy than the global alpha model. (**B**) The norm of the model weights is compared between the two models. Because the local alpha model optimizes the regularization hyperparameter independently for each vertex, the norm of the model weights can dramatically vary in scale across vertices. By contrast, the global alpha model optimizes a single regularization hyperparameter across all vertices, and thus the norm of the model weights is similar in scale across vertices. Because MC uses the Euclidean distance between model weights, the differences in scale across vertices in the local alpha model bias the recovered networks. (**C**) The weights of the local alpha model are rescaled according to model prediction accuracy in the training set, and the norm of the rescaled weights is compared to the norm of the weights of the global alpha model. Rescaling the weights of the local alpha model produces weights with norms that are comparable to the norms obtained with the global alpha model, thus eliminating the biases caused by differences in the scale of the model weights when fit with local alphas.
